# Comparative analysis of molecular representations in prediction of drug combination effects

**DOI:** 10.1101/2021.04.16.439299

**Authors:** B. Zagidullin, Z. Wang, Y. Guan, E. Pitkänen, J. Tang

## Abstract

Application of machine and deep learning methods in drug discovery and cancer research has gained a considerable amount of attention in the past years. As the field grows, it becomes crucial to systematically evaluate the performance of novel computational solutions in relation to established techniques. To this end we compare rule-based and data-driven molecular representations in prediction of drug combination sensitivity and drug synergy scores using standardized results of 14 throughput screening studies, comprising 64 200 unique combinations of 4 153 molecules tested in 112 cancer cell lines. We evaluate the clustering performance of molecular representations and quantify their similarity by adapting the Centered Kernel Alignment metric. Our work demonstrates that to identify an optimal molecular representation type it is necessary to supplement quantitative benchmark results with qualitative considerations, such as model interpretability and robustness, which may vary between and throughout preclinical drug development projects.

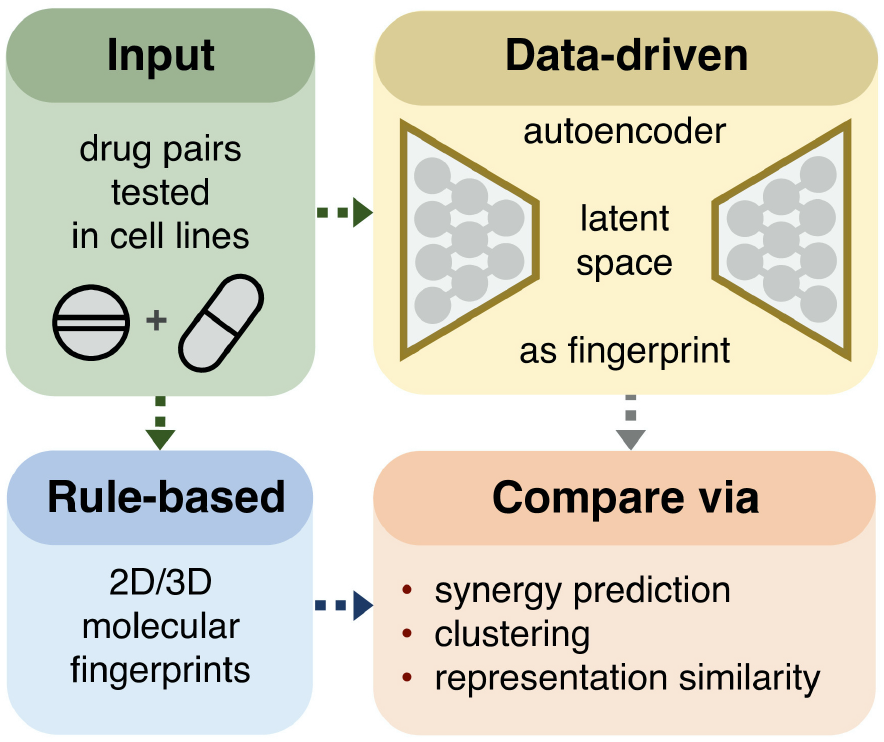

## 1 Introduction

In the past years deep learning (DL) methods have been successfully applied to a variety of research topics in biomedicine and drug discovery [1–3]. Deep neural networks achieve state-of-the-art performance in medical computer vision tasks and protein structural modelling, enabling de novo generation of drug candidates and development of prognostic clinical models [4–8]. However, such performance of DL models is context-dependent [9–12]. While quantitative metrics are routinely and effectively used to compare various computational methods, overreliance on them is a well-known issue [13–18]. It is beneficial to supplement performance results on benchmark datasets with estimates of model uncertainty and robustness, as well as ability to generalize on unseen data [19–21]. These aspects are particularly important in the biomedical research, where *in silico* model predictions direct experimental design choices, as exhaustively testing all combinations of relevant factors is usually unfeasible due to the combinatorial explosion [22,23].

Advances in high throughput screening of bioactive compounds in cancer cell lines promote the development of personalized cancer treatments [24]. A major goal in such drug sensitivity and resistance testing studies is to prioritize promising combinatorial therapies that involve coadministration of multiple drugs [25]. By combining synergistic compounds, often with distinct mechanisms of action, it is possible to overcome single-drug resistance, produce sustained clinical remissions and diminish adverse reactions [26,27]. Drug synergy refers to a degree of drug-drug interaction quantified as the difference between expected and observed dose-response profiles measured by a biological endpoint, such as cell viability or cell toxicity [28]. While synergy characterizes how compounds modulate each other’s biological activity, combination sensitivity score (CSS) quantifies drug combination efficacy [29]. In addition to the CSS, we use four synergy scores based on distinct null models, namely Bliss independence, Highest single agent, Loewe additivity, and Zero Interaction potency in the regression analysis of molecular fingerprints [30–34]. Predicting drug combination synergy and sensitivity is related to Quantitative Structure Activity Relationship (QSAR) modelling and Virtual Screening [35,36]. The QSAR captures mathematical associations between drug descriptors and assay endpoints based on the assumption that structurally similar compounds have similar bioactivity properties, while in the Virtual Screening studies candidate molecules are prioritized for subsequent experimental validation according to in silico prediction results [37]. Rule-based molecular fingerprints are commonly used as drug descriptors in QSAR/Virtual Screening, and MACCS structural keys based on molecular topology are arguably the most popular type of rule-based fingerprints [38–41]. Other types include circular topological fingerprints that describe combinations of non-hydrogen atom types and paths between them within a predefined atom neighborhood, and pharmacophore fingerprints that incorporate local features related to molecular recognition [42-44].

More recently, data-driven fingerprints generated by DL models have been shown to perform well in various research projects [45]. Majority of such DL fingerprints are based on the encoder-decoder architecture, whereby an approximate identity function is learned to translate high-dimensional input into a low-dimensional, fixed-size latent manifold, which is then used to reconstruct the original input [46]. When an encoder-decoder DL model is trained on chemical structures, its latent manifold is interpreted as a data-driven fingerprint. Examples of early DL fingerprinting models include a Convolutional Neural Network (CNN), Chemception, and a Recurrent Neural Network, SMILES2Vec, as well as a Variational Autoencoder model with a CNN encoder and a Gated Recurrent Unit-based decoder [47–51]. Development of attention methods for sequence modelling further contributed to the popularity of data-driven DL fingerprints, whereas evolution of generative models enabled de novo molecular design through latent space sampling [52–56]. These DL solutions operate on images of molecules or SMARTS/SMILES sequences to create drug structural representations [57–59]. Further, DL fingerprints may be enriched with numerical drug descriptors through multitask DL learning methods or simply by concatenating to latent space [60]. Unlike sequence-based versions, Geometric Deep Learning fingerprints are derived from molecular graphs, and in addition to global molecular descriptors enable position-aware encoding of individual atom and bond features [61–67].

There exist several extensive benchmark datasets for ranking DL models in chemoinformatics tasks, such as MoleculeNet, Open Graph Benchmark and Benchmarking GNNs [68–70]. Despite the widespread use of molecular fingerprints, there is a lack of systematic evaluation of data-driven DL and rule-based versions. To address the gap, we study 11 types of molecular representations, comprising seven DL and four rule-based variants, in prediction of cancer drug combination synergy and sensitivity, based on 17 271 848 data points from 14 cancer drug screening studies (Fig.1, experiment VS I). By comparing four synergy scores based on distinct null models we identify a preferred synergy formulation for use in cancer drug combinations research [71,72]. We measure the fingerprint similarity by adapting Centered Kernel Alignment as a distance metric (Fig.1, experiment VS II). Lastly, we explore the downstream performance of molecular representations by clustering compounds assigned to 10 Anatomical Therapeutic Chemical (ATC) classes in one-vs-all mode (Fig.1, experiment VS III). We believe that our work will contribute to the rational design of drug combinations, enable easier selection of molecular representations for in silico modelling, and promote further use of Deep Learning methods in biomedicine.

**Figure 1:**
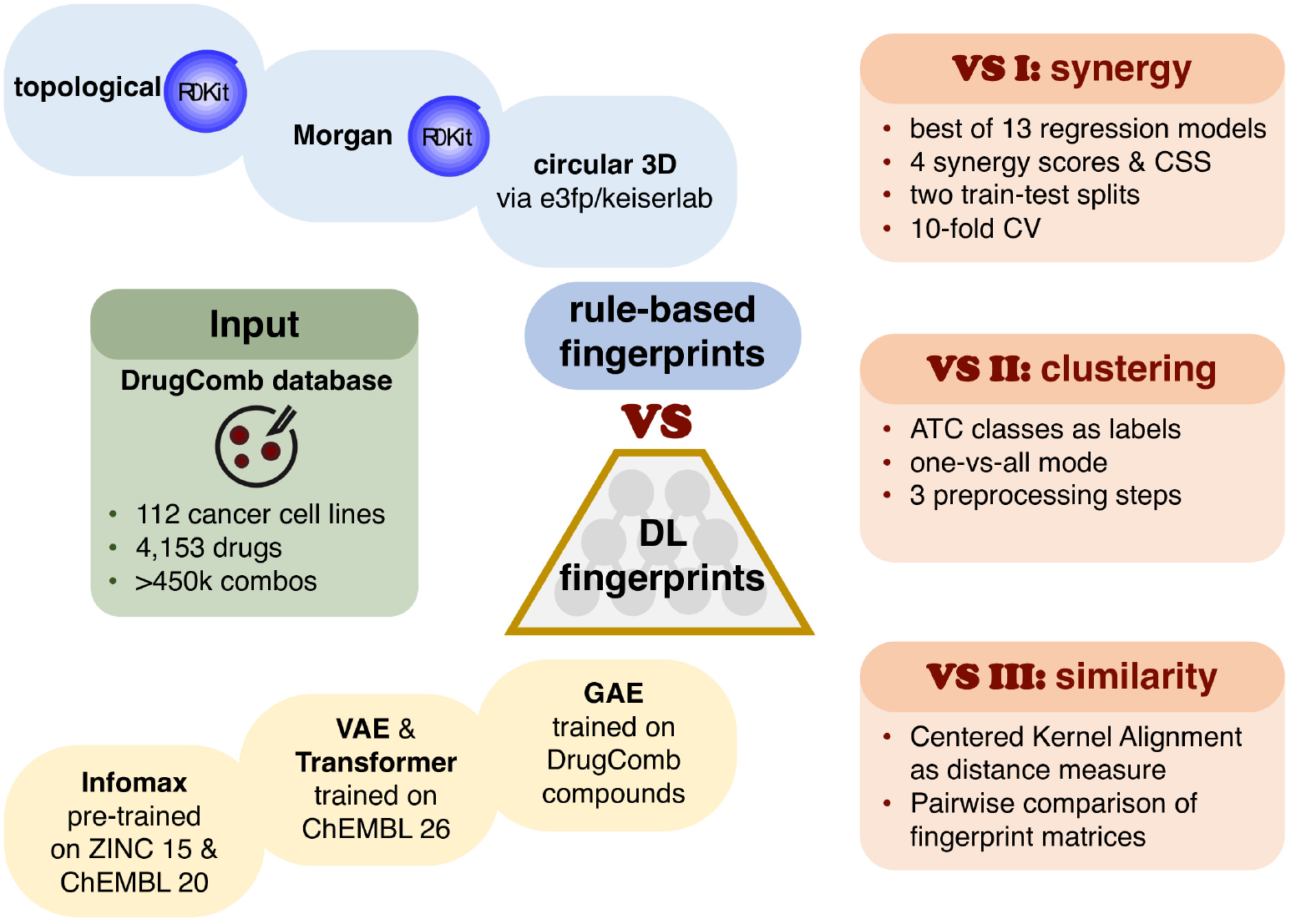
Study workflow. Compounds found in combinations in the DrugComb database are represented using four rule-based (blue) and seven data-driven (yellow) fingerprint types. Rule-based fingerprints include topological, and 2D/3D extended connectivity variants. Data-driven fingerprints are generated using two Variational Autoencoders and two Transformer models trained on ChEMBL 26, Graph Autoencoder trained on DrugComb compounds, and a pre-trained Deep Graph Infomax model (Infomax). The fingerprints are compared in three tasks: predictions of drug combination sensitivity and four synergy scores (**VS I**); representation similarity based on Centered Kernel Alignment (**VS II**); one-vs-all fingerprint clustering based on ATC drug classes (**VS III**). **VS I** results are also used to identify the most predictive synergy model.

## 2 Methods

### 2.1 Data provenance

The DrugComb data portal, one of the largest public drug combination databases, is used to access combination sensitivity and synergy data [73]. Its October 2 019 release contains standardized and harmonized results of 14 drug sensitivity and resistance studies on 4 153 drug-like compounds screened in 112 cell lines for a total of 447 993 drug-drug-cell line tuples. Each pairwise drug combination is characterized by the combination sensitivity score (CSS) and four synergy scores, namely Bliss independence (Bliss), Highest single agent (HSA), Loewe additivity (Loewe), and Zero interaction potency (ZIP), further details are in the Supplementary Information. ChEMBL (release 26) is used to obtain SMILES strings, which are subsequently standardized by stripping salt residues and solvent molecules [74,75]. SMILES shorter than 8 and longer than 140 characters are filtered out. PubChem identifiers (CID) are used to cross-reference compounds between the databases when necessary. The final DL training dataset consists of 1 795 483 unique SMILES with a median length of 48 and a median absolute deviation of 10.

### 2.2 Molecular representations

Fingerprints are numeric arrays of *n* elements (bits) long, where n ranges between 16 and 1 024 depending on fingerprint type. Even though n values up to 16,384 have been tested in literature demonstrating a positive correlation between fingerprint size and downstream prediction performance, not all the studies support these findings [38,76]. Fingerprints used in the current work are classified into rule-based with binary values and Deep learning-based with continuous values. Rule-based models are further split into topological, 2D and 3D circular subtypes. Deep Learning fingerprints are split into sequence and graph subtypes. More detailed classification is found in Table 1.

**Table 1:**
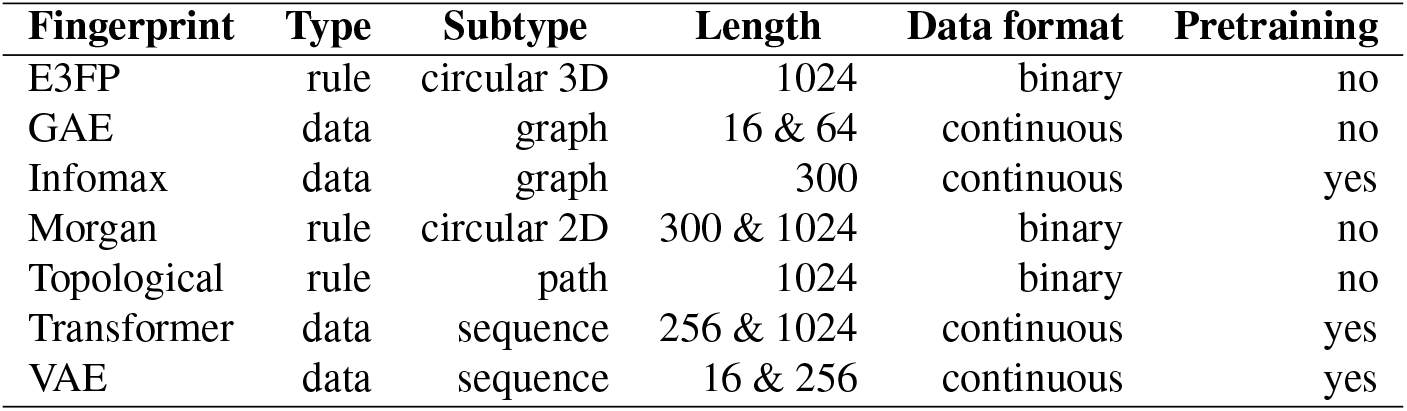
Pearson correlation coefficients of 10 regression algorithms in prediction of synergy and sensitivity scores based on Infomax 300 and Morgan 1 024 bits long fingerprints with one-hot encoded cell line labels as inputs. Averaged results of three independent 5-fold CV rounds on 10% of data run with models’ default hyperparameters.

#### 2.2.1 Rule-based fingerprints

Four types of rule-based fingerprints used in the current work are: path-based (Topological 1 024 bits longs), 2D circular (Morgan 300 and 1 024 bits long) and 3D circular (E3FP 1 024 bits). Topological and Morgan variants are selected due to their good performance in Virtual Screening experiments [38,43]. E3FP is a 3D extension of 2D extended-connectivity models, it is generated following the no_Stereo variant [77].

#### 2.2.2 Deep learning-based fingerprints

Seven data-driven molecular fingerprints of different lengths are generated using four types of unsupervised encoder-decoder DL models, namely a Graph Autoencoder (GAE), a Variational Autoencoder (VAE), a Transformer, and a pre-trained Deep Graph Infomax (Infomax).

##### GAE fingerprints

16 bits long GAE fingerprints are defined via a diagonal semidefinite matrix of singular values Σ, obtained through the Singular Value Decomposition of GAE embedding matrix [61,62]. Inspired by the Ky Fan matrix k-norm, equal to the sum of k largest singular values of the matrix, the main diagonal of Σ is used as a 16 bits long fingerprint [78]. If small molecules result in diagonal shorter than 16 bits, then zero-padding is applied. 64 bits long GAE fingerprints are generated by concatenating average, min- and max-pooled representations of the embedding matrix to 16 bits long GAE fingerprints.

##### VAE fingerprints

VAE fingerprints are 16 and 256 bits long latent spaces of two independently trained VAE models [49].

##### Transformer fingerprints

64 bits long Transformer fingerprints are constructed by concatenating average- and max-pooled latent embeddings of the 16 bits model with the first output of its last and second last recurrent layers. Similarly, the 1 024 bits variant is generated from the embedding space of the Transformer 256 bits model [52].

##### Infomax fingerprints

Infomax fingerprints are 300 bits long, generated using a pre-trained Deep Graph Infomax model that by design maximizes mutual information between local and global molecular graph features [79,80].

### 2.3 Deep Learning models used for fingerprint generation

#### Graph Autoencoder model

Graph Autoencoder model uses a graph 𝒢, with 𝒱 nodes and ε edges as input, where 𝒱 correspond to atoms and ε to atomic bonds. Additional numeric features may be incorporated via node or bond feature matrices [81]. A graph 𝒢 is represented with an adjacency matrix, A *ϵ* ℝ^|𝒱|×|𝒱|^, where |𝒱 |are node indices, such that non-zero A elements correspond to existing molecular bonds [82]. A is normalized to be symmetric and contain self-loops following [83]:

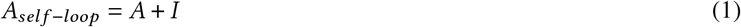

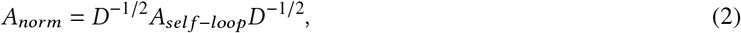

where *I* is an identity matrix equal in size to *A, D* is a diagonal node degree matrix such that its main diagonal represents bond counts of *A*_*self* −*loop*_. GAE model is initialized with a node matrix of 54 atom features, where each atom is represented by an array of one-hot encoded values denoting one of the 37 atoms types, six possible atom degrees, five atom charges, four variants of chiral configuration and an aromaticity indicator, all generated using RDKit. One-hot refers to encoding categorical variables as binary arrays. To make the GAE model compatible with previously unseen atoms a placeholder for an unknown atom type is added. GAE encoder consists of seven convolutional layers with sum pooling followed by ReLu activation [84]. The decoder part is a dot product of the embedding matrix with itself, followed by 0.1 dropout and sigmoid activation. Cross-entropy over *A*_*norm*_ is used as a loss function. Empty nodes in *A*_*norm*_ are initialized with zeros.

#### Variational Autoencoder model

Two VAE models are trained with the embedding sizes of 16 or 256 bits. Both models have a 54 characters in vocabulary, consisting of 53 unique alphanumeric characters found in SMILES and an additional empty character for zero-padding. Input length is 140 characters, zero-padded if necessary. VAE encoder consists of three 1D convolutional layers of 9, 9 and 10 neurons, each followed by SELU activation [85]. The decoder consists of three GRU layers with a hidden dimension of 501, followed by softmax activation. Loss function is an equally weighted combination of binary cross-entropy and Kullback–Leibler divergence. Xavier uniform initialization is used to assign the starting weights of two VAE models [86].

#### Transformer model

Two Transformers are trained with the embedding sizes of 16 or 256 bits. The vocabulary size for both models is 58 characters including 53 unique SMILES characters and five tokens for *end-of-string, mask, zero-pad, unknown-character* and *initialize-decoding*. Maximum input length is 141 characters, zero-padded if necessary. Both models have four-headed attention and six transformer layers, with a dropout of 0.3 applied to the positional encoding output [54]. Loss function is negative log likelihood. Network weights are initialized with Xavier uniform.

#### Deep Graph Infomax (Infomax) model

Infomax is pre-trained on 465 000 molecules from ChEMBL 20 and on 2 million molecules from ZINC 15 by Hu *et*.*al*. [80,87].

#### DL Model training

The GAE model is trained on 4 153 DrugComb drug-like compounds, while VAE and Transformer models are trained on 1 795 483 molecules from DrugComb and ChEMBL 26 databases. Five-fold cross-validation is used for training all the DL models. Transformer and VAE models are trained for 10 epochs on each fold, GAE is trained for 40 epochs on each fold. All models use Adam optimizer with a learning rate decay and an initial learning rate of 1 · 10^−3^, the training is halted once the learning rate reaches 1 · 10^−6^ or loss reaches zero [70,88–90]. GAE hyperparameters are optimized using tree-structured parzen estimators with a budget of 1 000 iterations, other DL models employ random search [91]. Further training details can be found in *Supplementary Information*.

### 2.4 Regression analysis of molecular fingerprints - VS I

#### Data input

One-hot encoded cell line labels and each of 11 drug fingerprints are used as inputs to regression models to predict drug combinations sensitivity and synergy. Combination fingerprints are generated by concatenating single molecular representations, topological fingerprints are bit-averaged [92]. Full dataset contains 362 635 cell line-drug combination tuples of 3 421 compounds, when filtered by the SMILES strings (SMILES-filtered), and 447 993 combination tuples of 4 153 molecules, when filtered by the CID (CID-filtered). For each cell line-drug combination tuple, four synergy scores and CSS sensitivity scores are obtained from DrugComb. If found, biological replicates are averaged, further, dose-dependent synergy scores are averaged inside cell line-drug combination tuple.

#### Model selection

Model selection for the regression analysis of molecular fingerprints is split into three steps. In the first step, 13 different regression models are tested thrice in five-fold cross-validation on the 10% of the full dataset, sampled without replacement (*Supplementary Information*). The goal is to identify an optimal type of a regression model for prediction of four synergy scores and the CSS. The second step concerns hyperparameter tuning of the previously selected regression model on all available data in ten-fold cross-validation. Lastly, the model is trained in ten-fold repeated cross-validation on SMILES-filtered and CID-filtered datasets with 90:10 and 60:40 train:test splits [93].

#### Regression performance metrics

Pearson Correlation Coefficient (PCC) and Root Mean Squared Error (RMSE) are used to assess the regression performance. PCC and RMSE 95% confidence intervals are calculated via Student’s t-distribution estimate of Fisher’s z transformed PCC values, and via empirical bootstrap with 1 000 iterations and symmetric confidence intervals [94–98]. RMSE values are normalized by standard deviations. Shapiro-Wilk test is used to test the normality assumption [99].

#### Related work

PCC scores of regression models, used to predict single synergy scores in three recent studies, are in the *Supplementary Information* section.

### 2.5 Fingerprint Similarity - VS II and III

#### Fingerprint similarity metric - VS II

All 11 types of molecular representations vary in length and data types, making commonly-used metrics, such as Jaccard–Tanimoto or cosine distances poor choices for fingerprint comparison [100–102]. Jaccard–Tanimoto is suboptimal, as it is based on bits present in one fingerprint, absent in another, and shared by both [103]. Cosine distance between two vectors, defined as their inner product normalized by the corresponding L2 norms, only measures an angle between two vectors without accounting for differences in their ranges [104]. It may be possible to post-process DL fingerprints and define common distance metrics on both the binary and real-valued arrays [105]. However, we opted against it, as we are not aware of any studies that systematically assess DL fingerprint similarity or quantify downstream effects of such transformations. Recall that an inner product is an unnormalized measure of similarity allowing metrics based on the Canonical Correlation Analysis (CCA), singular vector CCA and projection-weighted CCA to be defined on any real-valued arrays [106–108]. All these methods underperform when the number of compounds is smaller than the dimensionality of feature space, *i*.*e. n* bits [109]. It is not intuitive to use unnormalized inner product as a similarity measure, as it is unbounded and requires original data to be referenced alongside the similarity scores. Since calculation of pairwise compound distances is not a prerequisite to quantify their similarity, we compare complete fingerprint matrices using Centered Kernel Alignment (CKA), a modification of Hilbert-Schmidt Independence Criterion (HSIC) originally proposed to assess nonlinear dependence of two sets of variables [110].

#### Fingerprint matrix - VS II and III

Let *m* compounds be represented with two fingerprint matrices *X* and *Y*, where individual fingerprints *x*_*i*_ and *y*_*i*_ may be of different lengths x and y and data types:

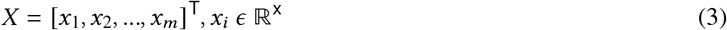

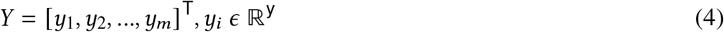

*X* and *Y* are normalized by subtracting column means from the corresponding column values.

#### Linear kernel k - VS II

Let *K* be a kernel matrix, such that its entries correspond to scalar outputs of a linear kernel function k. Let k be an inner product, 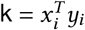, where *x*_*i*_ and *y*_*i*_ are 1D vectors from two fingerprint matrices *X* and *Y* corresponding to the same compound or feature. When *x*_*i*_ and *y*_*i*_ are column vectors, *K* becomes a feature similarity matrix:

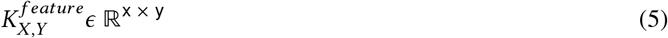

If *x*_*i*_ and *y*_*i*_ are row vectors, *K* is a sample similarity matrix:

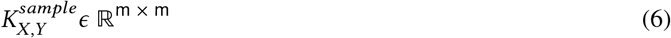

#### Hilbert-Schmidt Independence Criterion - VS II

Hilbert-Schmidt Independence Criterion (HSIC) is a test statistic equal to 0 when *X* and *Y* are independent [103]. Unnormalized HSIC is without an upper bound and equal to:

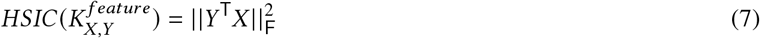

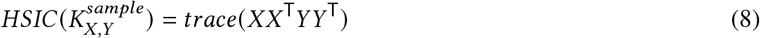

where *Y* ^T^*X* is a dot product of feature vectors and 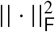 is a squared Frobenius norm and *XX*^T^ and *YY*^T^ are left Gram matrices. Notice that:

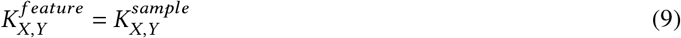

Further, for centered *X* and *Y* under linear dot product kernel:

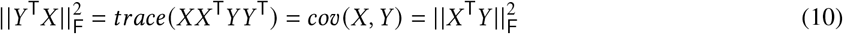

where *cov* (*X, Y*) is a cross-covariance matrix of *X* and *Y* [104].

#### Centered Kernel Alignment - VS II

HSIC is an empirical statistic that converges to its unbiased value at a rate of 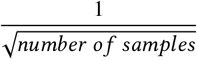 [105]. Unbiased HSIC values are used to define Centered Kernel Alignment (CKA), a normalized *number o f samples* version of HSIC that ranges from 0 to 1. CKA is used to quantify the difference between two fingerprint matrices *X* and *Y*. When CKA is calculated via the feature similarity, it is defined as:

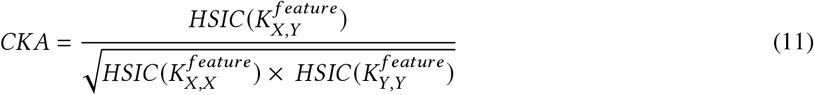

CKA is a non-linear extension of the CCA and does not require any assumptions about noise distributions in the datasets [112]. CKA with linear kernel is equivalent to the RV coefficient and Tucker’s Congruence coefficient [109,113–115]. If the number of samples is higher than the number of features, CKA should be calculated using feature similarities. Conversely, sample space and use of Gram matrices is preferred.

#### Fingerprint clustering - VS III

The Anatomical Therapeutic Chemical (ATC) Classification System is used to annotate drugs according to biological systems on which they act, as well as their therapeutic, pharmacological, and chemical properties [116]. The 2 228 DrugComb compounds found in the ATC database are assigned to 10 classes. All but GAE 16 bits and Morgan 1 024 bits models are then used to generate nine fingerprint matrices. The generated fingerprint matrices are preprocessed three-fold: by z-score normalization, z-score normalization followed by dimensionality reduction with PCA, and z-score normalization followed by dimensionality reduction with PLS. For the PCA preprocessing the number of loadings explaining >0.95 variance is used, PLS regression for dimensionality reduction is performed with ATC labels as targets. Linear Discriminant Analysis (LDA) is used for one-vs-all clustering with ATC class labels as response variables, averaged Silhouette score and Variance Ratio Criterion are clustering performance metrics [117,118].

#### Silhouette score - VS III

Silhouette for a single point is defined as:

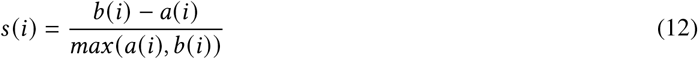

where *a* is the mean distance between point *i* and all points within its cluster *C*_*i*_ and *b* is the smallest mean distance between point *i* and all points in a cluster ≠ *C*_*i*_.

#### Variance Ratio Criterion - VS III

Variance Ratio Criterion (VRC) is a ratio of between-to within-cluster variation, adjusted by the number of clusters. VRC is closely related to the F-statistic in ANOVA [119]. Both scores are min-max scaled to be in [0, 1].

### 2.6 Computational facilities

All models are trained on Tesla P100 PCIe 16GB GPU. VM deployment is automated with Docker 19.03.9, python-openstack 3.14.13 and Heat Orchestration Template, Newton release. All experiments are performed using: Catboost 0.24, DGL 0.5, numpy 1.19.1, mlxtend 0.17.3, Optuna 2.2.0, pandas 1.1.3, Python 3.7.6, PyTorch 1.6, RDKit 2020.03.2, scikit-learn 0.23.2, scipy 1.4.1, XGBoost 1.2.1. Figures are created in R ggplot2 3.3.2, matplotlib 3.3.2 and seaborn 0.11.

## 3 Results and Discussion

### 3.1 Prediction of drug combination synergy and sensitivity - VS I

#### Regression model selection

We identify Catboost Gradient Boosting on Decision Trees (GBDT) as an optimal regression model for the prediction of drug combination sensitivity and synergy after testing 13 algorithms on the 10% of the DrugComb dataset in three replicates (Table 2). Three of the tested algorithms failed to generate any predictions and are omitted. With optimized hyperparameters GBDT models tend to reach the early stopping criterion in the last 20% of the training on all the fingerprint variants which indicates correctly tuned hyperparameters, further details are in Supplementary Information. There exist alternative dataset splitting modes that incorporate chemical similarity via Tanimoto distance or Murcko decomposition [68,120]. While they may better mimic current drug development practices and lead to a better correlation between in silico predictions and prospective experimental validation, we do not expect them to produce categorically different results.

**Table 2:**
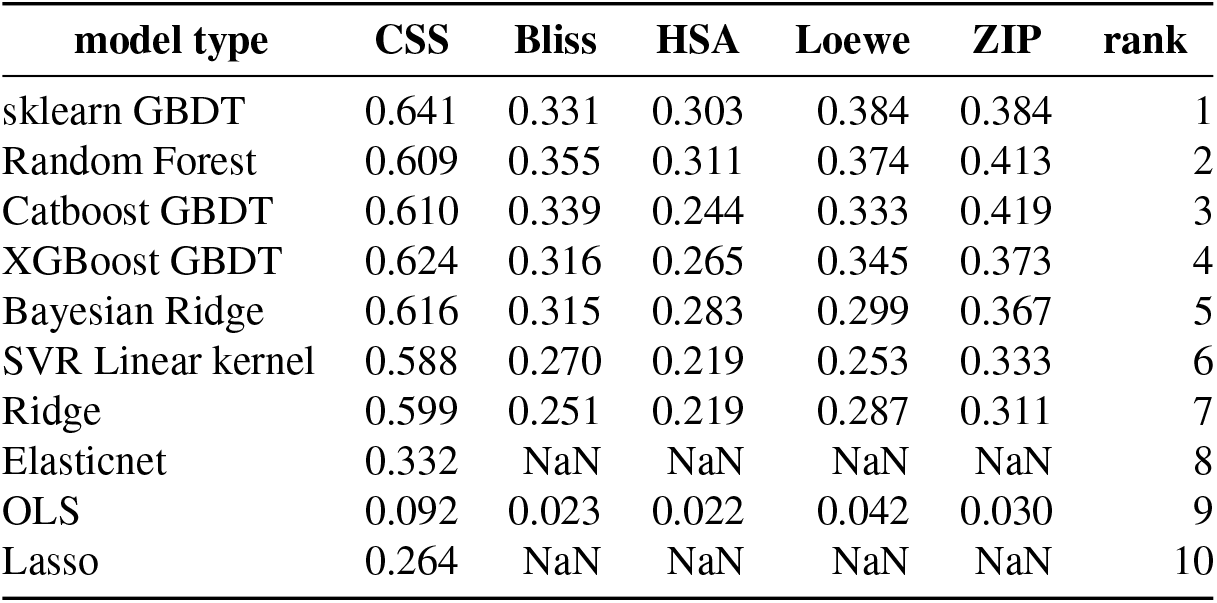
Pearson correlation coefficients of 10 regression algorithms in prediction of synergy and sensitivity scores based on Infomax 300 and Morgan 1 024 bits long fingerprints with one-hot encoded cell line labels as inputs. Models are trained in three replicates, with default hyperparameters in five-fold CV on 10% of data. VS I

| model type | CSS | Bliss | HSA | Loewe | ZIP | rank |
| --- | --- | --- | --- | --- | --- | --- |
| sklearn GBDT | 0.641 | 0.331 | 0.303 | 0.384 | 0.384 | 1 |
| Random Forest | 0.609 | 0.355 | 0.311 | 0.374 | 0.413 | 2 |
| Catboost GBDT | 0.610 | 0.339 | 0.244 | 0.333 | 0.419 | 3 |
| XGBoost GBDT | 0.624 | 0.316 | 0.265 | 0.345 | 0.373 | 4 |
| Bayesian Ridge | 0.616 | 0.315 | 0.283 | 0.299 | 0.367 | 5 |
| SVR Linear kernel | 0.588 | 0.270 | 0.219 | 0.253 | 0.333 | 6 |
| Ridge | 0.599 | 0.251 | 0.219 | 0.287 | 0.311 | 7 |
| Elasticnet | 0.332 | NaN | NaN | NaN | NaN | 8 |
| OLS | 0.092 | 0.023 | 0.022 | 0.042 | 0.030 | 9 |
| Lasso | 0.264 | NaN | NaN | NaN | NaN | 10 |

#### Regression performance

Among 11 fingerprinting models, Infomax 300 and VAE 256 achieved the highest PCC in prediction of Loewe synergy score using Catboost Gradient Boosting across all the test folds, cross-validation modes and duplicate filtering methods. As seen in Figure 2 and Table 3, for the 60 : 40 splits on the SMILES-filtered dataset Infomax reaches a PCC of 0.6842, while VAE 256 score is 0.6813. All tested fingerprints result in the CSS prediction scores above 0.85 PCC, with Infomax 300 and VAE 256 fingerprints still ranked on top. Infomax 300 and VAE 256 have overlapping 95% confidence intervals, as such they are considered to be equally performant. E3FP is the best rule-based fingerprint and is among the top three in most experimental runs. As seen in Figure 3 and Table 4, normalized RMSE scores further corroborate that DL-based fingerprints are better than rule-based variants in regression tasks. Further regression results for 90 : 10 and 60 : 40 train : test splits using SMILES and CID-filtered datasets are in the *Supplementary Figures and Tables* section (Figures S1-S6 and Tables S1-S6).

**Figure 2:**
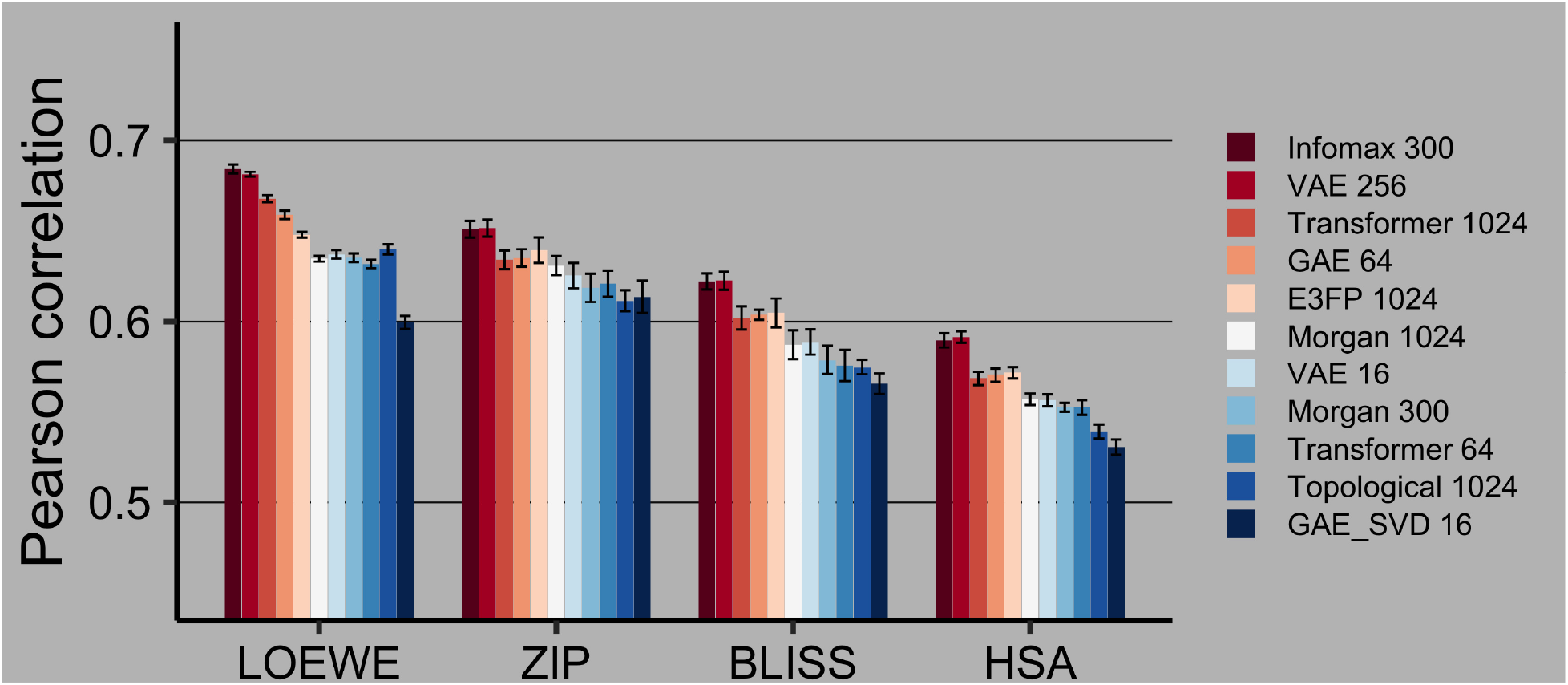
Drug combination synergy prediction on the SMILES-filtered dataset in 60 : 40 train:test split. 95% confidence intervals are calculated via Fisher z-transformation. Best models are highlighted with red. VS I

**Table 3:**
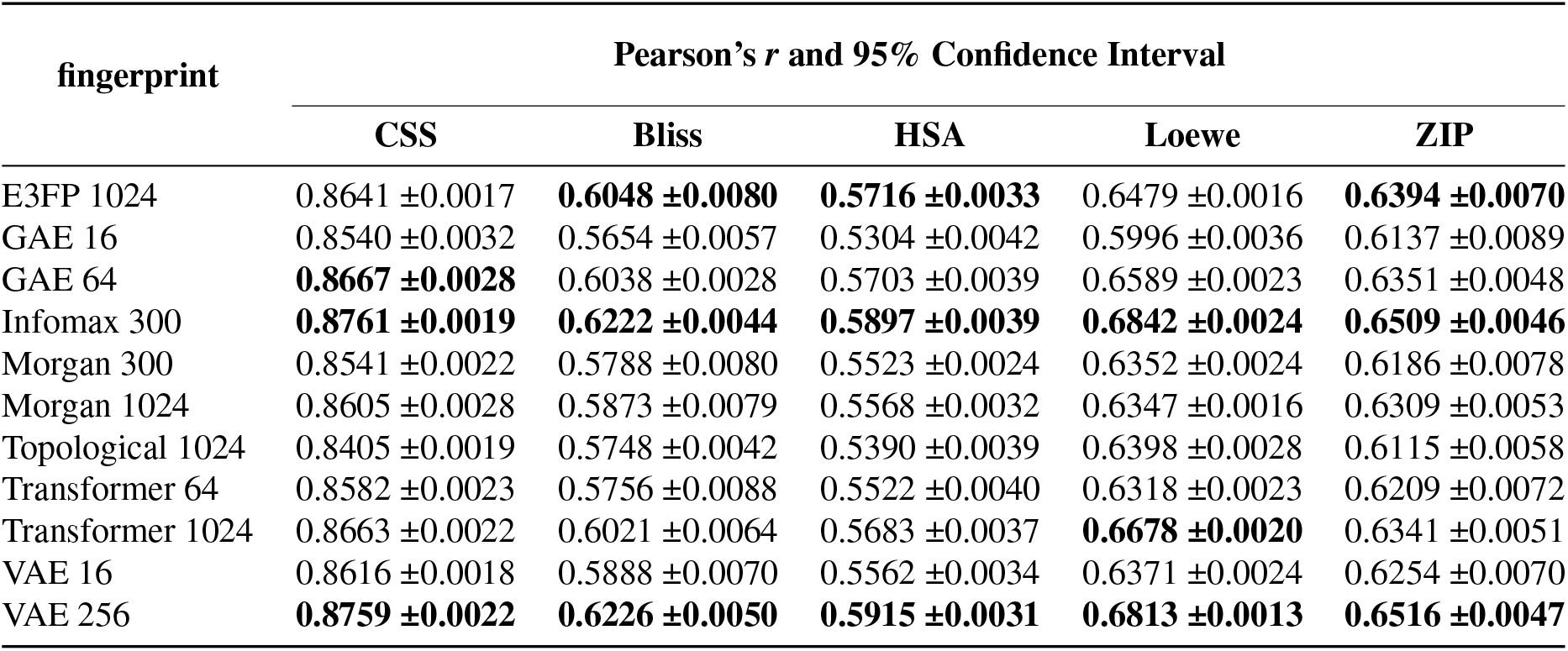
Drug combination synergy prediction on the SMILES-filtered dataset in 60 : 40 train:test split. 95% confidence intervals are calculated via Fisher z-transformation. Best models are highlighted with bold. VS I

| fingerprint | Pearson’s $r$ and 95% Confidence Interval | | | | |
| --- | --- | --- | --- | --- | --- |
|  | CSS | Bliss | HSA | Loewe | ZIP |
| E3FP 1024 | 0.8641 ±0.0017 | **0.6048 ±0.0080** | **0.5716 ±0.0033** | 0.6479 ±0.0016 | **0.6394 ±0.0070** |
| GAE 16 | 0.8540 ±0.0032 | 0.5654 ±0.0057 | 0.5304 ±0.0042 | 0.5996 ±0.0036 | 0.6137 ±0.0089 |
| GAE 64 | **0.8667 ±0.0028** | 0.6038 ±0.0028 | 0.5703 ±0.0039 | 0.6589 ±0.0023 | 0.6351 ±0.0048 |
| Infomax 300 | **0.8761 ±0.0019** | **0.6222 ±0.0044** | **0.5897 ±0.0039** | **0.6842 ±0.0024** | **0.6509 ±0.0046** |
| Morgan 300 | 0.8541 ±0.0022 | 0.5788 ±0.0080 | 0.5523 ±0.0024 | 0.6352 ±0.0024 | 0.6186 ±0.0078 |
| Morgan 1024 | 0.8605 ±0.0028 | 0.5873 ±0.0079 | 0.5568 ±0.0032 | 0.6347 ±0.0016 | 0.6309 ±0.0053 |
| Topological 1024 | 0.8405 ±0.0019 | 0.5748 ±0.0042 | 0.5390 ±0.0039 | 0.6398 ±0.0028 | 0.6115 ±0.0058 |
| Transformer 64 | 0.8582 ±0.0023 | 0.5756 ±0.0088 | 0.5522 ±0.0040 | 0.6318 ±0.0023 | 0.6209 ±0.0072 |
| Transformer 1024 | 0.8663 ±0.0022 | 0.6021 ±0.0064 | 0.5683 ±0.0037 | **0.6678 ±0.0020** | 0.6341 ±0.0051 |
| VAE 16 | 0.8616 ±0.0018 | 0.5888 ±0.0070 | 0.5562 ±0.0034 | 0.6371 ±0.0024 | 0.6254 ±0.0070 |
| VAE 256 | **0.8759 ±0.0022** | **0.6226 ±0.0050** | **0.5915 ±0.0031** | **0.6813 ±0.0013** | **0.6516 ±0.0047** |

**Figure 3:**
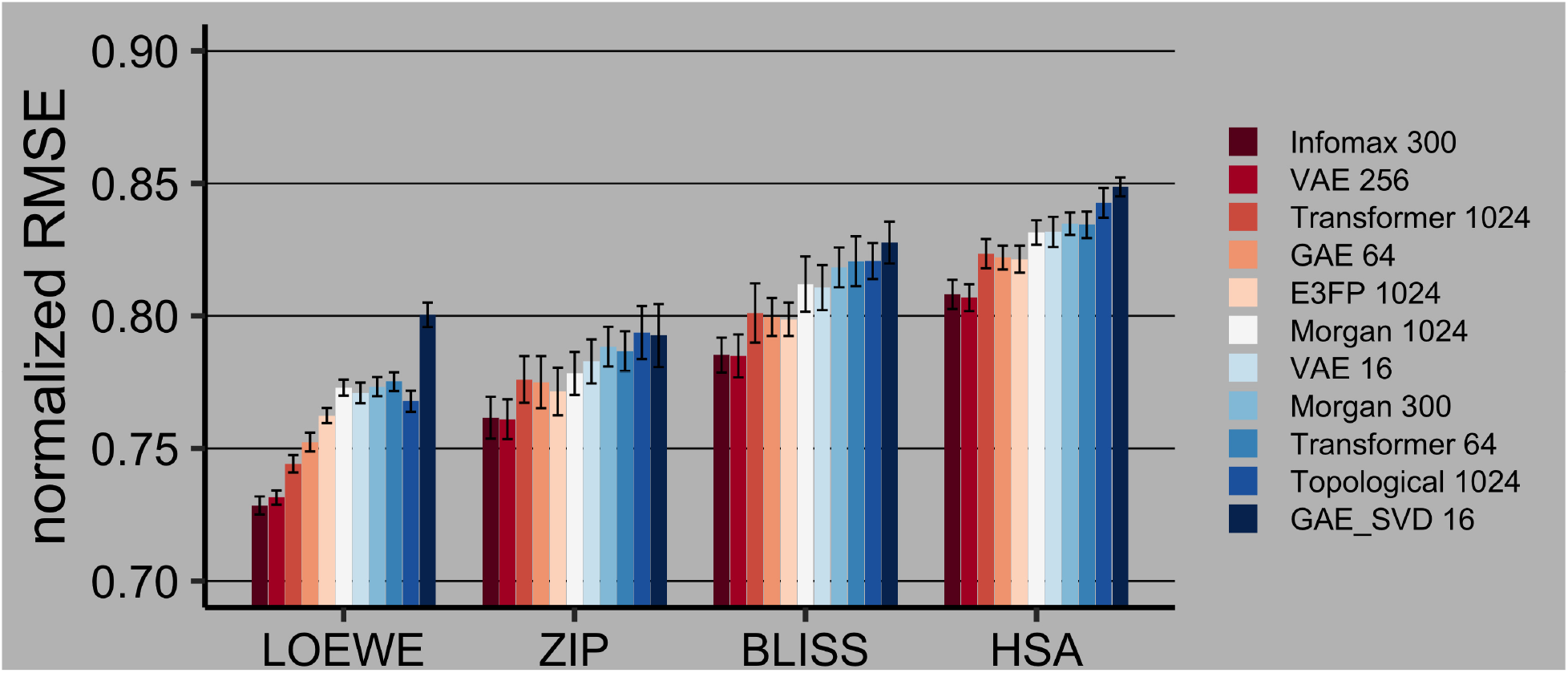
Drug combination synergy prediction on the SMILES-filtered dataset in 60 : 40 train:test split using RMSE, normalized by the target’s standard deviation. 95% confidence intervals are calculated via empirical bootstrap. Best models are highlighted with red. Normalized RMSE value of 1 indicates that the standard deviation of residuals is equal to the standard deviation of the target, *i*.*e*. a model that predicts mean values for all targets would have such a normalized RMSE. VS I

**Table 4:** Drug combination sensitivity and synergy prediction on the SMILES-filtered dataset in 60 : 40 train:test split using RMSE, normalized by the target’s standard deviation. 95% confidence intervals are calculated via empirical bootstrap. Best models are in bold. VS I

| fingerprint | normalized Root Mean Squared Error and 95% Confidence Interval |  |  |  |  |
| --- | --- | --- | --- | --- | --- |
|  | CSS | Bliss | HSA | Loewe | ZIP |
| E3FP 1024 | 0.5034 ± 0.0011 | **0.7987 ± 0.0063** | **0.8214 ± 0.0051** | 0.7624 ± 0.0029 | **0.7715 ± 0.0090** |
| GAE 16 | 0.5205 ± 0.0020 | 0.8277 ± 0.0079 | 0.8487 ± 0.0036 | 0.8004 ± 0.0046 | 0.7926 ± 0.0119 |
| GAE 64 | **0.4990 ± 0.0016** | 0.7996 ± 0.0072 | 0.8220 ± 0.0045 | 0.7524 ± 0.0035 | 0.7750 ± 0.0098 |
| Infomax 300 | **0.4822 ± 0.0011** | **0.7852 ± 0.0066** | **0.8081 ± 0.0055** | **0.7285 ± 0.0034** | **0.7616 ± 0.0079** |
| Morgan 300 | 0.5204 ± 0.0007 | 0.8183 ± 0.0075 | 0.8348 ± 0.0042 | 0.7733 ± 0.0036 | 0.7884 ± 0.0075 |
| Morgan 1024 | 0.5097 ± 0.0013 | 0.8120 ± 0.0104 | 0.8315 ± 0.0046 | 0.7729 ± 0.0030 | 0.7783 ± 0.0081 |
| Topological 1024 | 0.5420 ± 0.0011 | 0.8207 ± 0.0068 | 0.8427 ± 0.0056 | 0.7678 ± 0.0040 | 0.7937 ± 0.0100 |
| Transformer 64 | 0.5134 ± 0.0014 | 0.8206 ± 0.0094 | 0.8344 ± 0.0050 | 0.7752 ± 0.0035 | 0.7867 ± 0.0075 |
| Transformer 1024 | 0.4998 ± 0.0012 | 0.8011 ± 0.0112 | 0.8235 ± 0.0055 | **0.7442 ± 0.0033** | 0.7760 ± 0.0088 |
| VAE 16 | 0.5080 ± 0.0010 | 0.8107 ± 0.0085 | 0.8317 ± 0.0057 | 0.7709 ± 0.0039 | 0.7828 ± 0.0083 |
| VAE 256 | **0.4825 ± 0.0013** | **0.7849 ± 0.0081** | **0.8069 ± 0.0051** | **0.7315 ± 0.0027** | **0.7610 ± 0.0075** |

Experimental results indicate that if similarity-based clustering or identification of key molecular moieties are of interest, rule-based fingerprints should be considered. Their average performance in regression is compensated by the inbuilt interpretability and robust clustering performance [121]. On the other hand, neural fingerprint models are well-suited for regression tasks, as seen in the VS I experiment. It is important to note that the differences in regression performance between rule- and DL-based fingerprints do not exceed 0.05 PCC when predicting any synergy scores or the CSS. Consistently good performance of the DL models and E3FP fingerprints may be offset by their high computational costs during model training or fingerprint generation, respectively. GAE 64 fingerprints appear to be a reasonable compromise in terms of the downstream performance and relatively short model training times.

### 3.2 Fingerprint similarity - VS II and III

#### CKA distance (VS II)

A heatmap of pairwise CKA similarities between 11 fingerprints, as seen in Figure 4, indicates that similar types of fingerprints cluster together. Rule-based fingerprints form two clusters corresponding to topological and circular subtypes. All the DL fingerprints generated by the trained models form the third cluster. Graph-based models appear to be far removed from all sequence and rule-based variants. GAE 64 is the most different from other trained DL fingerprints, while being co-clustered with them. Infomax 300 fingerprints, based on a pre-trained Deep Graph Infomax model, are not part of any cluster. Smaller sequence-based DL fingerprints, namely VAE 16 and Transformer 64 are at least as similar to each other, as they are to their longer in-type/subtype counterparts. We conclude that fingerprint type and subtype, as indicated in Table 1, contribute the most to the CKA similarity, followed by fingerprint pretraining status, size, and data format.

**Figure 4:**
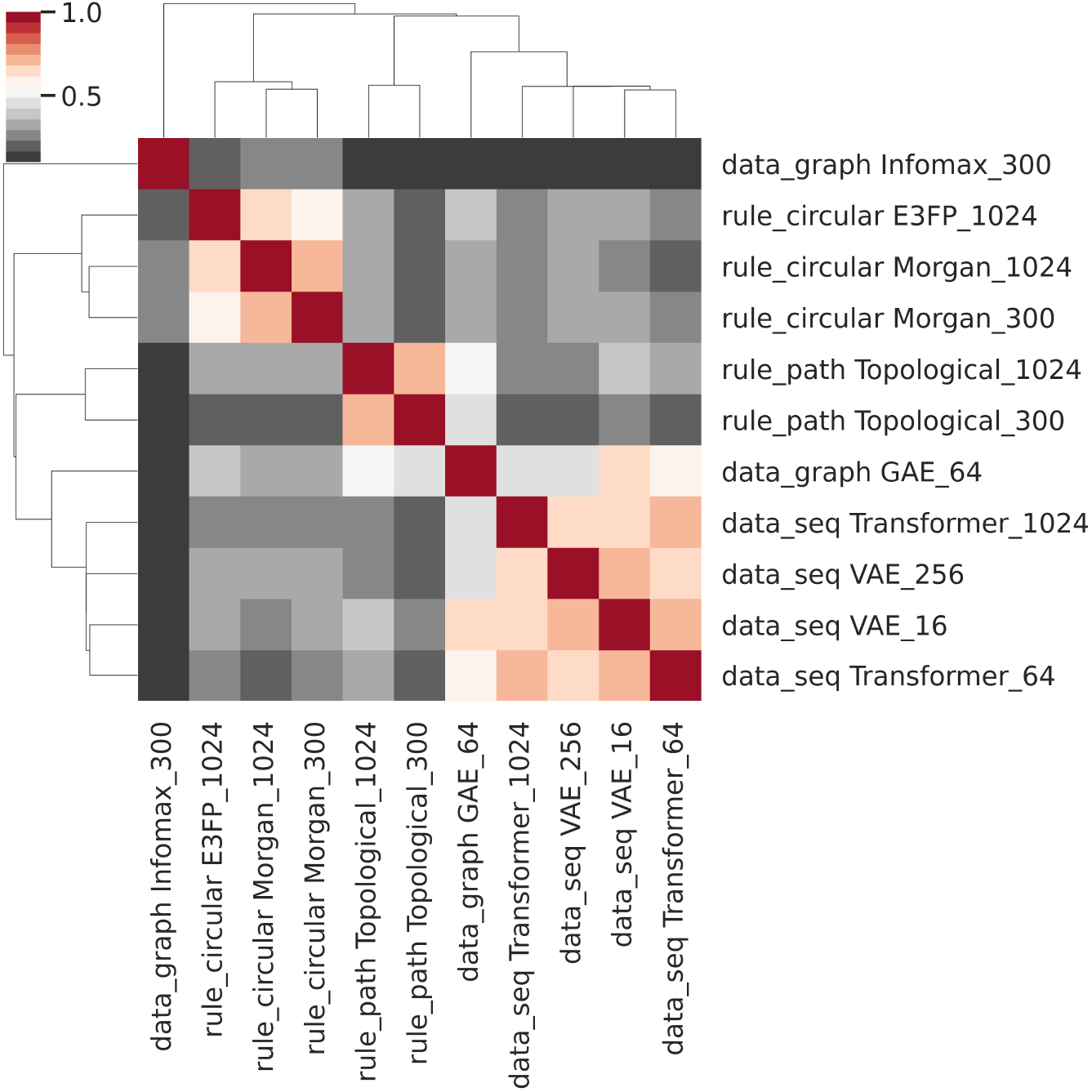
Heatmap of pairwise CKA distances between 11 fingerprints. Infomax and GAE 64 are Deep Learning fingerprints based on molecular graphs. VAE and Transformer are sequence-based DL fingerprints. E3FP and variants of Morgan and Topological fingerprints are generated using rule-based models. VS II

#### LDA clustering (VS III)

To further study the differences between fingerprint models, we perform one-vs-all LDA classification of 2 228 compounds based on their ATC classes, using nine different fingerprinting models to represent the molecules. The GAE 16 bits fingerprints are omitted, since GAE 64 bits fingerprints extend their shorter counterparts by concatenating average, min- and max-pooled embedding spaces. Further, due to the comparable performance of Morgan 300 and 1 024 bits models in VS I and VS II experiments, only Morgan 300 bits fingerprints are used in LDA clustering experiments. VS III clustering results are in Table 5 and the overview of DrugComb compounds with the corresponding ATC classes is in Figure 5. The Infomax 300 bits model achieves the best clustering results on the z-score normalized fingerprint matrices, followed by three rule-based fingerprints. Dimensionality reduction following z-score normalization generally improves clustering performance of all rule-based fingerprints. It has the opposite effect on most DL fingerprints, with the largest reduction seen in the Infomax 300 and GAE 64 models. Longer DL sequence models, namely VAE 256 and Transformer 1 024, perform better after dimensionality reduction steps, albeit with a minimal improvement in relative rankings. Such differences between graph and sequence-based DL fingerprints are supported by the CKA analysis (VS II), indicating that the graph-based fingerprints differ the most from other DL variants.

**Table 5:** One-vs-all LDA clustering in 10 ATC classes of 2 228 DrugComb compounds represented with nine fingerprint types. Averaged Silhouette and Variation Ratio Criterion (VRC) scores are rescaled to [0, 1]. Fingerprints are ranked according to scores on z-score normalized data. PCA and PLS-based dimensionality reduction improves rule-based fingerprint (denoted by R_ prefix) performance, most DL fingerprints (D_ prefix) decrease performance, VAE 256 and Transformer 1 024 benefit from dimensionality reduction, although minimally in terms of relative ranking. VS III

| fingerprint | z-score |  | z-score + PCA |  | z-score + PLS |  |
| --- | --- | --- | --- | --- | --- | --- |
|  | silhouette | VRC | silhouette | VRC | silhouette | VRC |
| D_Infomax 300 | 0.984 | 1.000 | 0.701 | 0.251 | 0.749 | 0.326 |
| R_E3FP 1024 | 1.000 | 0.831 | 0.993 | 0.958 | 0.988 | 0.959 |
| R_Topo 1024 | 0.979 | 0.771 | 1.000 | 0.997 | 1.000 | 1.000 |
| R_Morgan 300 | 0.967 | 0.655 | 0.993 | 1.000 | 0.997 | 0.984 |
| D_GAE 64 | 0.440 | 0.122 | 0.258 | 0.064 | 0.303 | 0.062 |
| D_VAE 256 | 0.399 | 0.049 | 0.547 | 0.170 | 0.574 | 0.190 |
| D_VAE 16 | 0.223 | 0.056 | 0.076 | 0.036 | 0.000 | 0.006 |
| D_Transformer 64 | 0.088 | 0.004 | 0.000 | 0.000 | 0.120 | 0.000 |
| D_Transformer 1024 | 0.000 | 0.000 | 0.131 | 0.023 | 0.377 | 0.067 |

**Figure 5:**
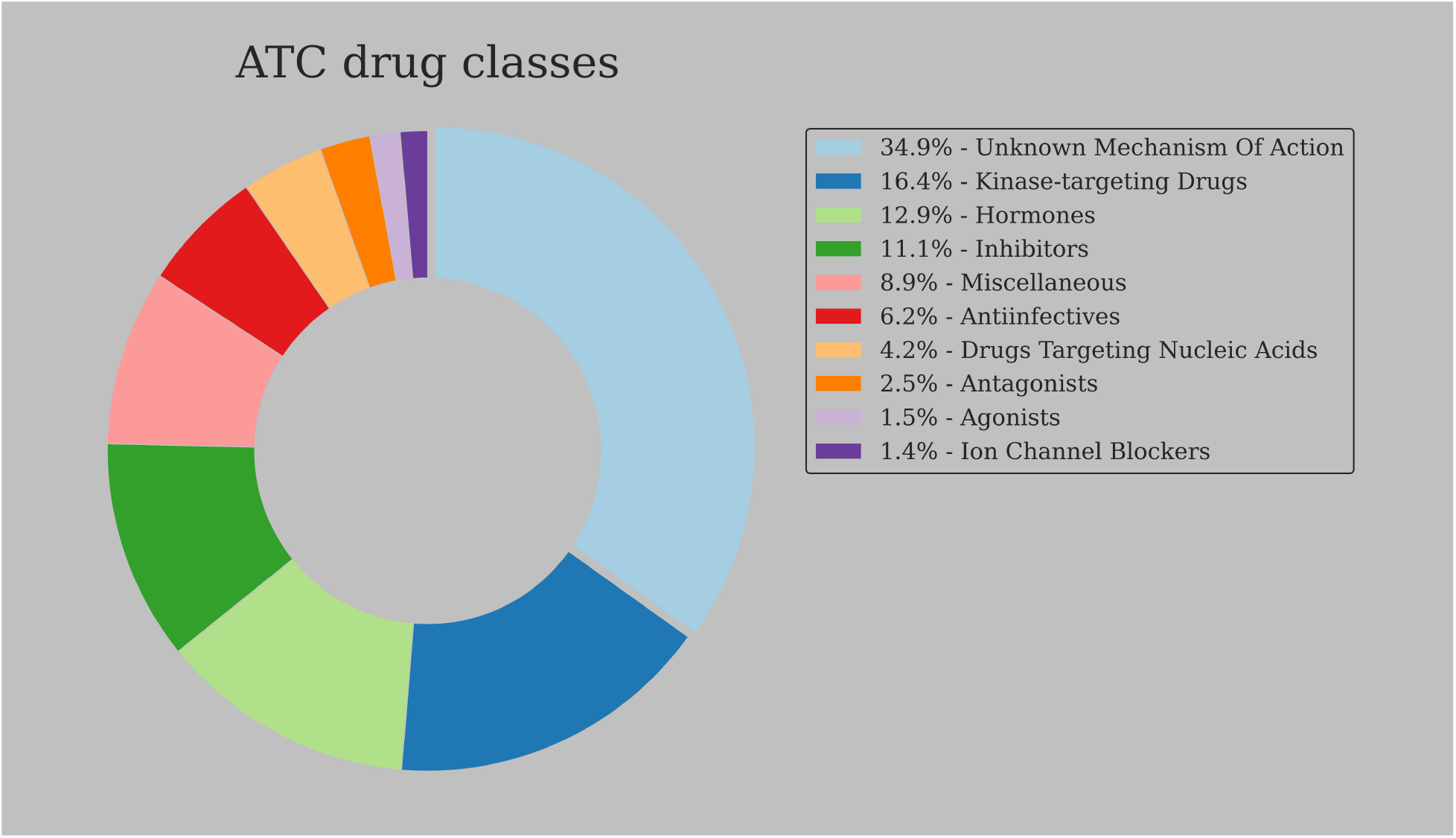
ATC drug classes of the DrugComb compounds (n=3 421). Over one third (n=1 193) compounds do not have a mechanism of action assigned in the ATC classification system. VS III

## 4 Conclusion

Choosing an optimal fingerprint type to represent molecular features is an important step in computational drug discovery. To this end, we systematically compared 11 variants of such molecular representations in predicting drug combination sensitivity and synergy scores, and evaluated their relationships based on the clustering performance and CKA-based fingerprint similarity. We found that VAE 256 bits long and 3D circular E3FP 1 024 bits long fingerprints generated from SMILES strings, as well as Infomax 300 bits long fingerprints based on molecular graphs lead to the best regression performance. Out of the four tested synergy scores, we observe that Loewe synergy is the easiest to predict with best models reaching PCC 0.72. CSS, a measure of drug combination efficacy, can be predicted >0.85 PCC with any fingerprint type. We found that the rule-based fingerprinting methods underperform in regression tasks in comparison to the data-driven DL variants. However, the gap between the best and worst performing fingerprint models rarely exceeds PCC 0.05. Further, we adapted Centered Kernel Alignment to quantify the extent of similarity between fingerprint matrices and to demonstrate that similar types of fingerprints cluster together. An optimal similarity measure for the comparison of single rule-based and data-driven fingerprints remains an open question. Lastly, in one-vs-all compound clustering using ATC classes as labels rule-based fingerprints perform on par or better than the best DL representations.

We conclude that the quantitative performance differences between rule-based and Deep Learning-based fingerprints are likely to be insignificant in the context of preclinical studies of small molecule drugs [122,123]. In order to identify an optimal fingerprint type for a given project we advise enriching quantitative performance metrics with qualitative concerns, e.g. available chemometric and deep learning expertise, model interpretability requirements, opinions of project stakeholders and model performance on unseen data. Fingerprints generated using the E3FP 1 024, Infomax 300, Morgan 1 024 and VAE 256 bits models are suggested as good starting points based on our experimental results and distinct methodologies underlying their data generating methods [124]. We recommend the Loewe synergy score for use in drug combination screening due to its best performance among four tested synergy models tested on dose-response data from 14 DSRT studies.

This work focuses on the evaluation of single fingerprint types. However, it is worth exploring the impact of combining several fingerprints together. We expect a statistically significant regression performance increase when combining molecular representations with low CKA similarity, or using models trained on multimodal data and/or key biological databases, such as Gene Ontology, Protein Data Bank and UniRef [5,125–127]. Another line of inquiry could address high computational costs of DL and E3FP models. To this end we suggest exploring alternative molecular representations and CPU-friendly generative models based on genetic algorithms, such as STONED on SELFIES [128]. Finally, we hope that in the future biomedical DL research will go beyond representation learning and will be used to derive novel biological knowledge by e.g. inferring synthetic and retrosynthetic chemical reactions, identifying novel disease-associated druggable proteins and clinically actionable biomarkers [129–131].

## 5 Key points

- To choose an optimal molecular fingerprint type it is advised to enrich quantitative metrics of model performance with qualitative concerns related to the nature of downstream tasks, model interpretability and robustness requirements, as well as available chemometric expertise.
- Data-driven fingerprints, namely VAE 256 bits long trained on SMILES and Infomax 300 bits long trained molecular graphs are well-suited for regression tasks. 1 024 bits long 2D and 3D circular fingerprints are flexible and well-performant rule-based models fit for clustering tasks. Graph Autoencoder 64 bits long model may be used in any analysis scenario as a baseline option.
- Loewe synergy scores enable the highest correlation between *in silico* predictions and subsequent experimental validation of drug combination synergy in cancer cell lines.
- Centered Kernel Alignment is an effective measure of molecular representation similarity applicable to any combination of rule-based and DL fingerprints.

## 6 Data and code availability

The data and code underlying this article are available in the article and in its online supplementary material. Code: https://github.com/NetPharMedGroup/publication_fingerprint/

Data: https://doi.org/10.5281/zenodo.4843919

## 7 Description of the authors

*Bulat Zagidullin* is a PhD candidate at the University of Helsinki. He obtained a BSc degree in Biochemical Engineering from Jacobs University Bremen and a MSc degree in Pharmaceutical Biotechnology from Martin Luther University Halle-Wittenberg.

*Ziyan Wang* is a Graduate student at the University of Michigan, Ann Arbor. He obtained a BSE in Computer Science from the University of Michigan.

*Yuanfang Guan* is an Associate Professor at the University of Michigan, Ann Arbor. She obtained a BSc in Biology in University of Hong Kong and a PhD in Molecular Biology at Princeton University.

*Esa Pitkänen* is a FIMM-EMBL Group Leader and an Academy of Finland Research Fellow at the Institute for Molecular Medicine Finland (FIMM), Helsinki. He received his PhD in Computer Science at the University of Helsinki.

*Jing Tang* is an Assistant Professor at the Faculty of Medicine, University of Helsinki. He obtained his PhD in Statistics from the University of Helsinki.

## 8 Acknowledgements

We acknowledge the computational resources provided by the Finnish IT Center for Science.

## 9 Funding

This work was supported by the European Research Council (ERC) starting grant DrugComb (Informatics approaches for the rational selection of personalized cancer drug combinations) [No. 716 063] (to BZ and JT). BZ is also supported by the faculty funded PhD position in the Integrative Life Science Doctoral Programme, University of Helsinki. EP is supported by the Academy of Finland grant [No. 322 675].

## 10 Potential conflicts of interest

None declared

## 11 Supplementary Information

### 11.1 Combination sensitivity and synergy scores

It is known that the choice of an appropriate null model of no interaction is crucial for an accurate assessment of drug synergy. Four such models are used in the current work. Bliss model is based on probabilistic independence of drug effects, such that single agents are competing, but independent perturbations each contributing to a total effect [30]. HSA assumes that an expected drug combination effect is the higher of the two single-agent effects at corresponding concentrations [31]. Degree of non-interaction according to Loewe is equal to the outcome of a sham experiment, that is combining a drug with itself [32,33]. Zero Interaction Potency (ZIP) assumes that non-interacting drugs minimally impact each other’s dose-response curves [34]. Combination Sensitivity Score quantifies drug combination efficacy, which is defined as an average of areas under drug combination dose-response curves, whereby each curve is determined by fixing one of the drugs at its IC50 concentration [29].

### 11.2 Regression model performance

13 regression models, representing a wide spectrum of ML algorithms, are compared in prediction of drug combination synergy and sensitivity. All models are tested with default hyperparameters in five-fold cross-validation using 1 024 bits long Morgan and 300 bits long Infomax fingerprints together with one-hot encoded cell line labels on 10% of randomly sampled data in three replicates. 13 tested regression models are: Bayesian Ridge, Catboost Gradient Boosting, ElasticNet, Gaussian Process Regression with a sum of Dot Product and White Kernels, Histogram-based Gradient Boosting, Isotonic Regression, Lasso regression, LassoLars regression, Linear regression, Ridge regression, Random Forest, Support Vector Machines with a linear kernel and XGBoost Gradient Boosting. All trees-based models are limited to a depth of six. Neural networks are not included in the comparison, since on tabular data they tend to perform on par with the previously mentioned methods, while being less interpretable and more difficult to set up [132,133].

Top four identified regression models are bagging and boosting ensembles, followed by linear kernel Support Vector Regression and Bayesian Ridge. Gaussian Process Regression, Isotonic and Lassolars models failed to generate any predictions and their results are omitted from Table 1. Catboost implementation of Gradient Boosting on Decision Trees (GBDT) is selected for further experiments due to its efficient GPU utilization and two design choices aimed at reducing overfitting: out-of-the-box categorical encoding that translates classes into numeric representations, binning them based on the expected value of target statistic, and Ordered Boosting whereby training data is randomly permuted throughout tree growing process to limit unwanted target metric prediction shift, one of well-known GBDT disadvantages [134–136].

Tuned in ten-fold cross-validation best hyperparameters for Catboost GBDT are Poisson bootstrap with 0.66 subsampling ratio, L2 regularization of 9, tree depth of 10, learning rate of 0.15 with 5 000 boosting iterations and 50 early stopping rounds. Overall, we conclude that ensembles are the most powerful type of tested ML algorithms in prediction of drug combination synergy and sensitivity [137].

### 11.3 Neural Network Training

For all DL models the hyperparameter search consisted of testing various activation functions (GELU, ELU, LeakyReLU, ReLU, SELU, Sigmoid, Softmax, Swish), dropout ratios (0.1 to 0.5 with 0.1 step size), initial learning rates (1 · 10^−1^ to 1 · 10^−5^, with a step size of 0.1), number of patience epochs (1 − 30 with a step size of 1) and learning rate decay factor (0.9 to 0.1 with a step size of 0.1). Both Transformer models were tested with 3− 6 attention heads. VAE encoders were tested with up to 5 convolutional layers and convolutional kernel sizes up to 10; VAE decoder is tested with up to 3 recurrent GRU layers of sizes up to 600.

#### Transformer

Two Transformer models are trained in 5-fold cross-validation, for up to 10 epochs on each fold with a decay factor of 0.1 and patience of 5 epochs in batch sizes of 650 and 340 for 16 bits and 256 bits long fingerprint variants. The final cross-entropy losses are 2 · 10^−7^ and 1 · 10^−8^, respectively.

#### VAE

Two Variational Autoencoder models with the latent spaces of 16 and 256 neurons are trained on ChEMBL 26 using 5-fold cross-validation and 10 epochs on each fold with a decay factor of 0.2 and patience of 2 and 3 epochs, respectively. An equally weighted sum of binary cross-entropy and KL-divergence is used as a loss metric. The final losses are 0.9231 and 0.0984 for 16 bits and 256 bits models.

#### GAE

A single Graph Autoencoder model is trained on 3 421 unique DrugComb compounds in 5-fold cross-validation mode for 200 epochs with a batch size of 340 for up to 40 epochs per fold. Learning rate decay factor is 0.1 with 30 epochs patience. Cross-entropy over the molecular graph adjacency matrix is used as a loss, with the best score of 0.8604 reached at the end of the 200^th^ epoch.

It is likely that longer training or a more extensive hyperparameter optimization or use of alternative optimizers, such as SGD with a cyclic learning rate scheduling, may result in lower final loss values, however, we do not expect it to have a significant influence on downstream experiments [138]. It is interesting to note that optuna-based optimization of the GAE model resulted in the encoder architecture consisting of seven convolutional layers 54 – 46− 40 −34 −28 −22 −16 neurons wide. Such a high number of convolutional layers is somewhat unexpected, as performance of Graph Neural Networks based on spectral convolutions is expected to deteriorate with the convolutional layer count above six, most likely due to excessive feature smoothing [139,140]. Such GAE encoder architecture may be explained by a positive correlation between the performance of Neural Networks and a number of trainable parameters, as a seven layer GAE model with dot-product based Decoder has *circa* 10k learnable parameters, whereas Transformer and VAE models have two orders of magnitude more [141].

### 11.4 Prior work in drug combination synergy predictions

In three independent studies listed below single synergy scores are predicted. Developed models are cross-validated on single datasets.

#### Random Forest

A Random Forest model developed by Menden *et*.*al*. achieved a 0.3 PCC in prediction of Loewe synergy using drug and cell line labels on the AstraZeneca dataset including 910 combinations tested in 85 cancer cell lines [142].

#### Convolutional Neural Network

A CNN model introduced by Preur *et*.*al*. achieved a 0.73 PCC in prediction of Loewe synergy from drug fingerprints, physicochemical molecular descriptors and basal expression of 3 984 cancer-associated genes on the O’Neil dataset including 583 drug combinations tested in 39 cancer cell lines [143,144].

#### Gradient Boosting and Random Forest

XGBoost and Random Forest models developed by Sidorov *et*.*al*. achieved a 0.64 PCC for a modified version of Bliss synergy from drug fingerprints and their physicochemical characteristics on the NCI60 Almanac dataset including 5 232 combinations tested in 60 cancer cell lines [26,145].

## 12 Supplementary Figures and Tables

**Figure S1:**
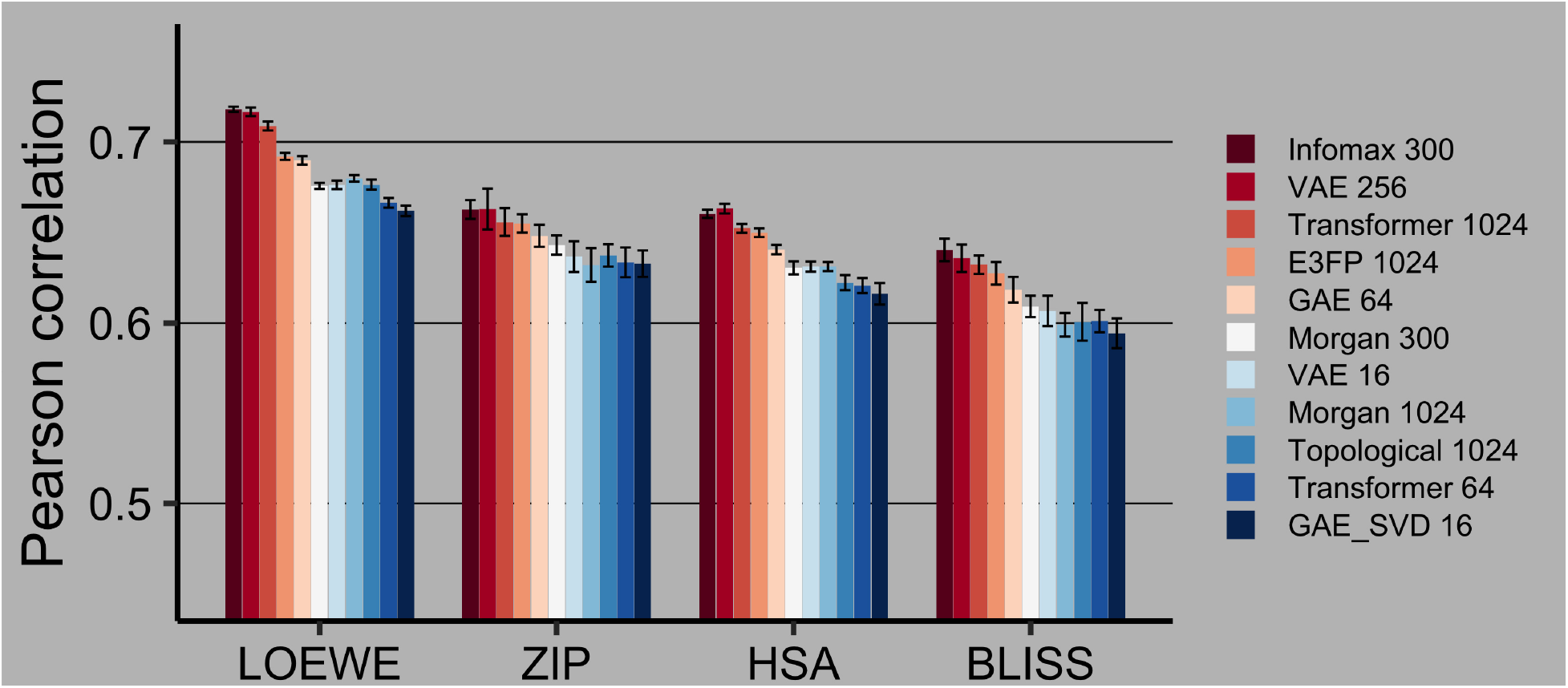
Drug combination synergy prediction on the CID-filtered dataset in 60 : 40 train:test split. 95% confidence intervals are calculated via Fisher z-transformation. Best models are highlighted with red. VS I

**Figure S2:**
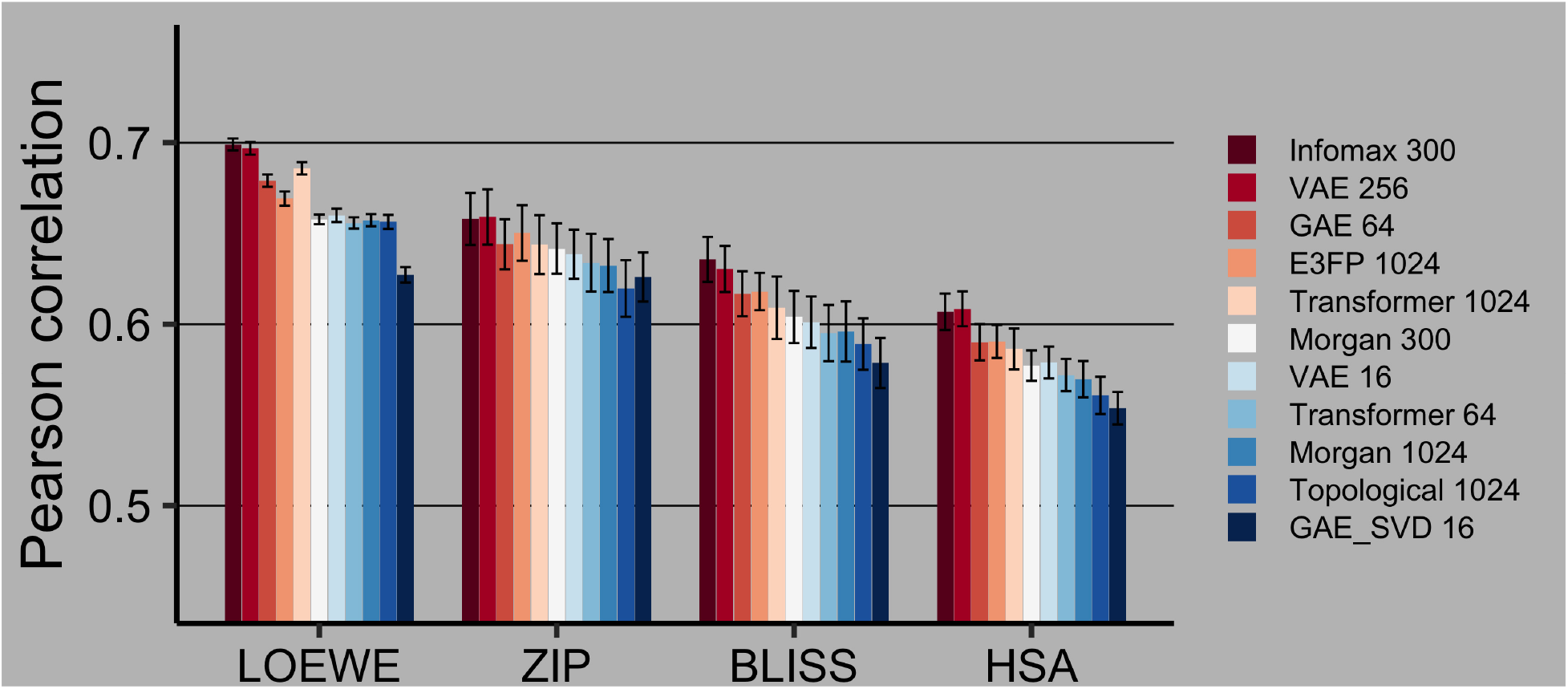
Drug combination synergy prediction on the SMILES-filtered dataset in 90 : 10 train:test split. 95% confidence intervals are calculated via Fisher z-transformation. Best models are highlighted with red. VS I

**Figure S3:**
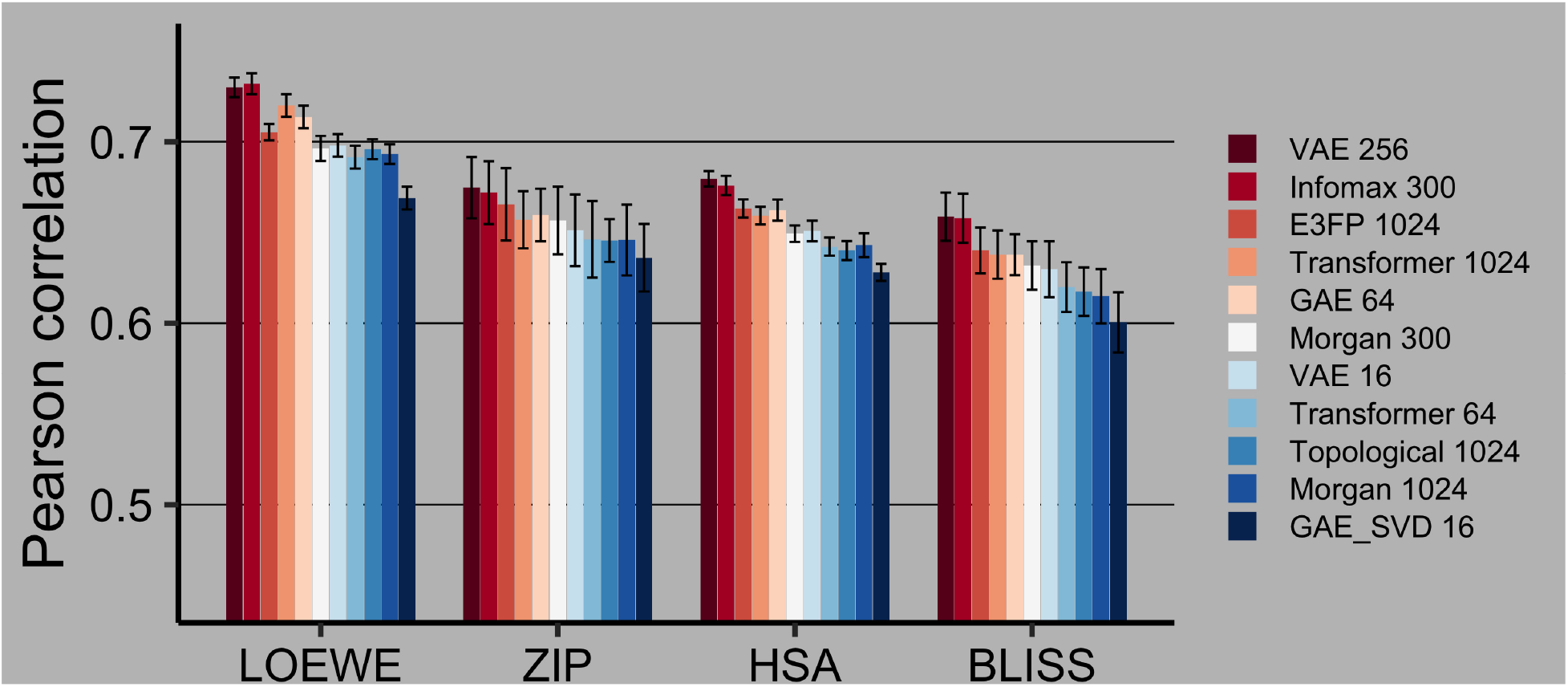
Drug combination synergy prediction on the CID-filtered dataset in 90 : 10 train:test split. 95% confidence intervals are calculated via Fisher z-transformation. Best models are highlighted with red. VS I

**Figure S4:**
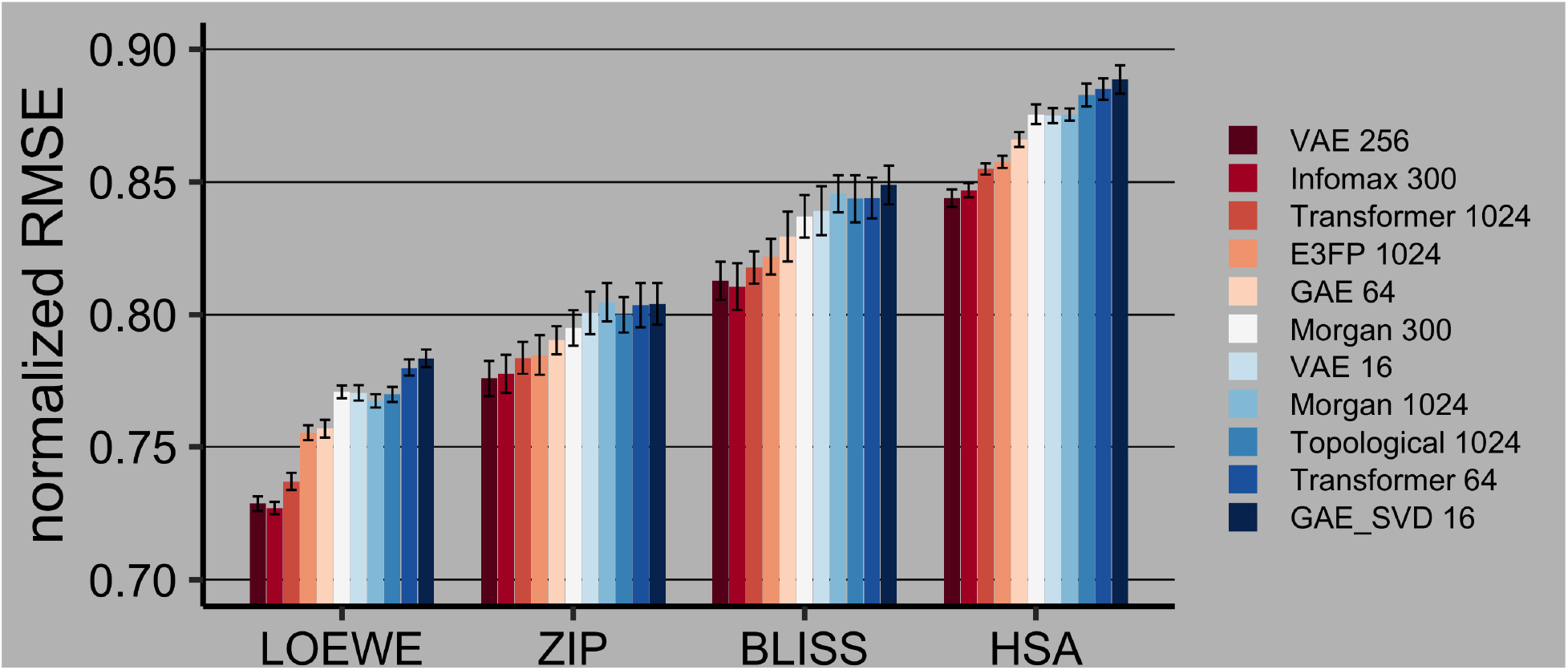
Drug combination synergy prediction on the CID-filtered dataset in 60 : 40 train:test split using RMSE, normalized by the target’s standard deviation. 95% confidence intervals are calculated via empirical bootstrap. Best models are highlighted with red. VS I

**Figure S5:**
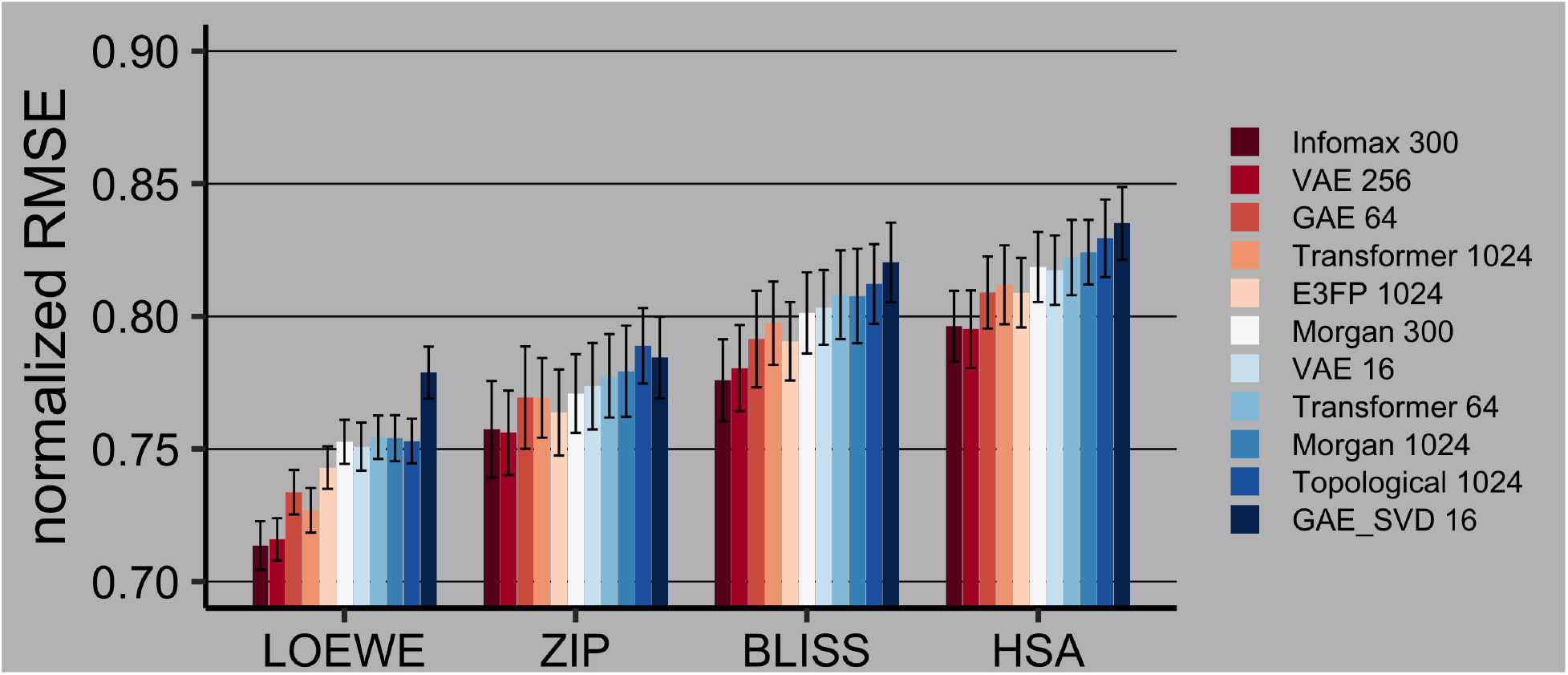
Drug combination synergy prediction on the SMILES-filtered dataset in 90 : 10 train:test split using RMSE, normalized by the target’s standard deviation. 95% confidence intervals are calculated via empirical bootstrap. Best models are highlighted with red. VS I

**Figure S6:**
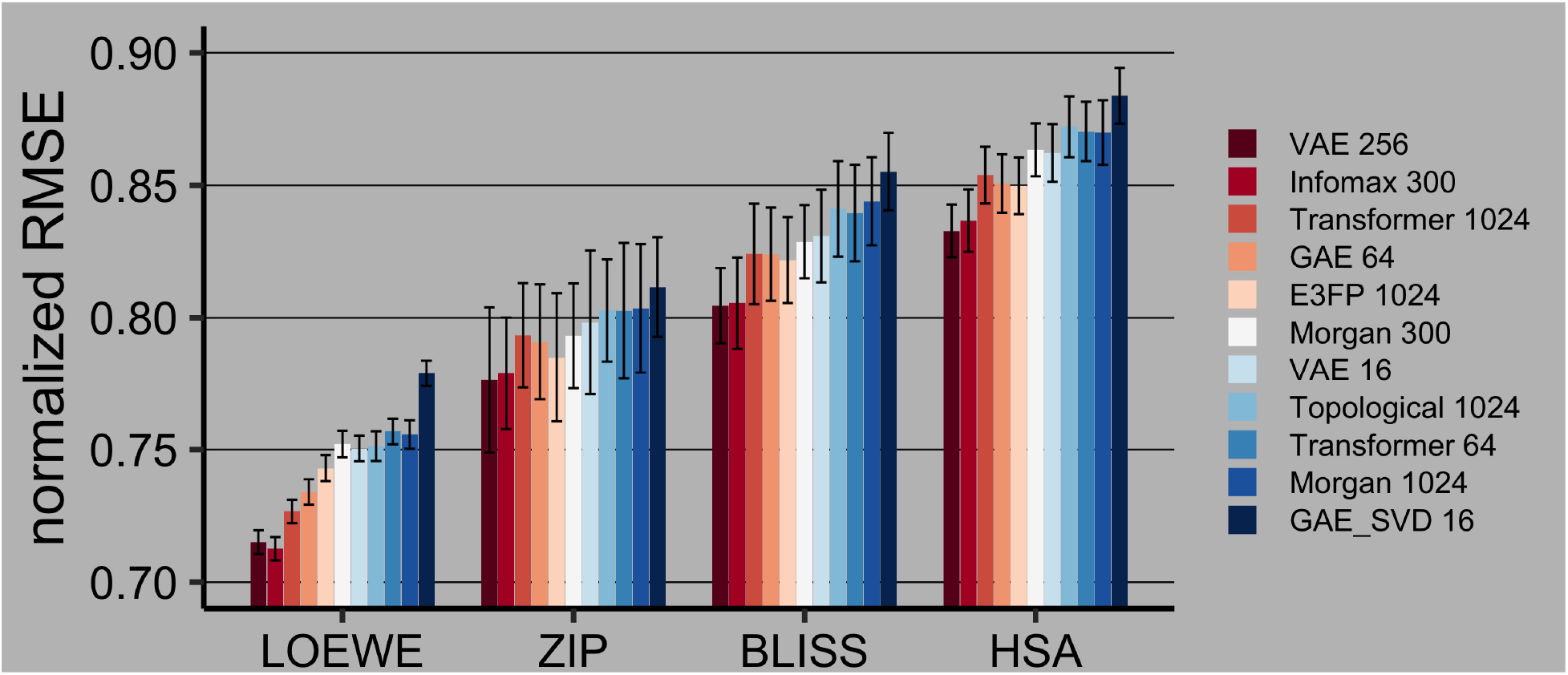
Drug combination synergy prediction on the CID-filtered dataset in 90 : 10 train:test split using RMSE, normalized by the target’s standard deviation. 95% confidence intervals are calculated via empirical bootstrap. Best models are highlighted with red. VS I

**Table S1:** Drug combination synergy prediction on the CID-filtered dataset in 60 : 40 train:test split. 95% confidence intervals are calculated via Fisher z-transformation. Best models are highlighted with bold. VS I

| fingerprint | Pearson's $r$ and 95% Confidence Interval | | | | |
| --- | --- | --- | --- | --- | --- |
|  | CSS | Bliss | HSA | Loewe | ZIP |
| E3FP 1024 | 0.8819 ±0.0019 | 0.6275 ±0.0062 | 0.6499 ±0.0024 | 0.6920 ±0.0019 | 0.6550 ±0.0051 |
| GAE 16 | 0.8744 ±0.0018 | 0.5944 ±0.0082 | 0.6162 ±0.0059 | 0.6619 ±0.0029 | 0.6328 ±0.0073 |
| GAE 64 | 0.8800 ±0.0022 | 0.6184 ±0.0071 | 0.6406 ±0.0020 | 0.6898 ±0.0023 | 0.6482 ±0.0061 |
| Infomax 300 | **0.8907 ±0.0016** | **0.6404 ±0.0062** | **0.6603 ±0.0022** | **0.7180 ±0.0014** | **0.6627 ±0.0052** |
| Morgan 300 | 0.8771 ±0.0017 | 0.6092 ±0.0059 | 0.6305 ±0.0036 | 0.6757 ±0.0017 | 0.6431 ±0.0053 |
| Morgan 1024 | 0.8712 ±0.0022 | 0.5990 ±0.0065 | 0.6312 ±0.0025 | 0.6799 ±0.0018 | 0.6321 ±0.0093 |
| Topological 1024 | 0.8718 ±0.0014 | 0.6007 ±0.0104 | 0.6223 ±0.0042 | 0.6764 ±0.0028 | 0.6374 ±0.0062 |
| Transformer 64 | 0.8716 ±0.0014 | 0.6011 ±0.0062 | 0.6207 ±0.0042 | 0.6664 ±0.0027 | 0.6335 ±0.0082 |
| Transformer 1024 | **0.8854 ±0.0016** | **0.6322 ±0.0051** | **0.6523 ±0.0025** | **0.7088 ±0.0025** | **0.6558 ±0.0077** |
| VAE 16 | 0.8767 ±0.0016 | 0.6067 ±0.0084 | 0.6312 ±0.0028 | 0.6763 ±0.0024 | 0.6367 ±0.0085 |
| VAE 256 | **0.8908 ±0.0023** | **0.6358 ±0.0076** | **0.6632 ±0.0027** | **0.7166 ±0.0024** | **0.6629 ±0.0113** |

**Table S2:** Drug combination synergy prediction on the SMILES-filtered dataset in 90 : 10 train:test split. 95% confidence intervals are calculated via Fisher z-transformation. Best models are highlighted with bold. VS I

| fingerprint | Pearson's $r$ and 95% Confidence Interval | | | | |
| --- | --- | --- | --- | --- | --- |
|  | CSS | Bliss | HSA | Loewe | ZIP |
| E3FP 1024 | 0.8732 ±0.0058 | **0.6180 ±0.0103** | **0.5905 ±0.0091** | 0.6692 ±0.0039 | **0.6503 ±0.0153** |
| GAE 16 | 0.8641 ±0.0063 | 0.5786 ±0.0138 | 0.5537 ±0.0090 | 0.6272 ±0.0043 | 0.6260 ±0.0135 |
| GAE 64 | **0.8754 ±0.0064** | 0.6168 ±0.0124 | 0.5901 ±0.0100 | 0.6791 ±0.0034 | **0.6440 ±0.0138** |
| Infomax 300 | **0.8826 ±0.0060** | **0.6357 ±0.0124** | **0.6069 ±0.0100** | **0.6990 ±0.0033** | **0.6580 ±0.0143** |
| Morgan 300 | 0.8702 ±0.0054 | 0.6040 ±0.0144 | 0.5772 ±0.0084 | 0.6578 ±0.0027 | 0.6417 ±0.0138 |
| Morgan 1024 | 0.8640 ±0.0052 | 0.5960 ±0.0166 | 0.5697 ±0.0100 | 0.6573 ±0.0034 | 0.6323 ±0.0146 |
| Topological 1024 | 0.8500 ±0.0044 | 0.5891 ±0.0141 | 0.5608 ±0.0103 | 0.6564 ±0.0039 | 0.6197 ±0.0156 |
| Transformer 64 | 0.8682 ±0.0047 | 0.5951 ±0.0155 | 0.5720 ±0.0089 | 0.6558 ±0.0031 | 0.6339 ±0.0159 |
| Transformer 1024 | 0.8742 ±0.0055 | 0.6091 ±0.0172 | 0.5864 ±0.0112 | **0.6859 ±0.0034** | 0.6439 ±0.0162 |
| VAE 16 | 0.8712 ±0.0062 | 0.6011 ±0.0142 | 0.5789 ±0.0088 | 0.6600 ±0.0037 | 0.6385 ±0.0135 |
| VAE 256 | **0.8829 ±0.0063** | **0.6304 ±0.0127** | **0.6085 ±0.0096** | **0.6969 ±0.0035** | **0.6591 ±0.0152** |

**Table S3:** Drug combination synergy prediction on the CID-filtered dataset in 90 : 10 train:test split. 95% confidence intervals are calculated via Fisher z-transformation. Best models are highlighted with bold. VS I

| fingerprint | Pearson's $r$ and 95% Confidence Interval | | | | |
| --- | --- | --- | --- | --- | --- |
|  | CSS | Bliss | HSA | Loewe | ZIP |
| E3FP 1024 | 0.8849 ±0.0049 | **0.6401 ±0.0126** | **0.6632 ±0.0050** | 0.7053 ±0.0045 | **0.6655 ±0.0199** |
| GAE 16 | 0.8761 ±0.0041 | 0.6005 ±0.0166 | 0.6280 ±0.0047 | 0.6690 ±0.0063 | 0.6361 ±0.0186 |
| GAE 64 | **0.8869 ±0.0050** | 0.6377 ±0.0113 | 0.6623 ±0.0058 | 0.7136 ±0.0062 | 0.6596 ±0.0145 |
| Infomax 300 | **0.8942 ±0.0056** | **0.6579 ±0.0135** | **0.6759 ±0.0053** | **0.7320 ±0.0058** | **0.6719 ±0.0173** |
| Morgan 300 | 0.8824 ±0.0045 | 0.6318 ±0.0133 | 0.6493 ±0.0045 | 0.6963 ±0.0069 | 0.6566 ±0.0187 |
| Morgan 1024 | 0.8748 ±0.0053 | 0.6149 ±0.0150 | 0.6430 ±0.0066 | 0.6932 ±0.0054 | 0.6458 ±0.0195 |
| TopoA 1024 | 0.8647 ±0.0049 | 0.6174 ±0.0133 | 0.6400 ±0.0052 | 0.6959 ±0.0055 | 0.6456 ±0.0118 |
| TB 64 | 0.8798 ±0.0059 | 0.6199 ±0.0137 | 0.6422 ±0.0050 | 0.6915 ±0.0063 | 0.6462 ±0.0211 |
| TB 1024 | **0.8853 ±0.0047** | 0.6378 ±0.0133 | 0.6593 ±0.0049 | **0.7200 ±0.0062** | 0.6570 ±0.0157 |
| VAE 16 | 0.8838 ±0.0053 | 0.6297 ±0.0154 | 0.6508 ±0.0057 | 0.6980 ±0.0062 | 0.6512 ±0.0197 |
| VAE 256 | 0.8949 ±0.0052 | **0.6587 ±0.0133** | **0.6796 ±0.0042** | **0.7300 ±0.0054** | **0.6747 ±0.0169** |

**Table S4:**
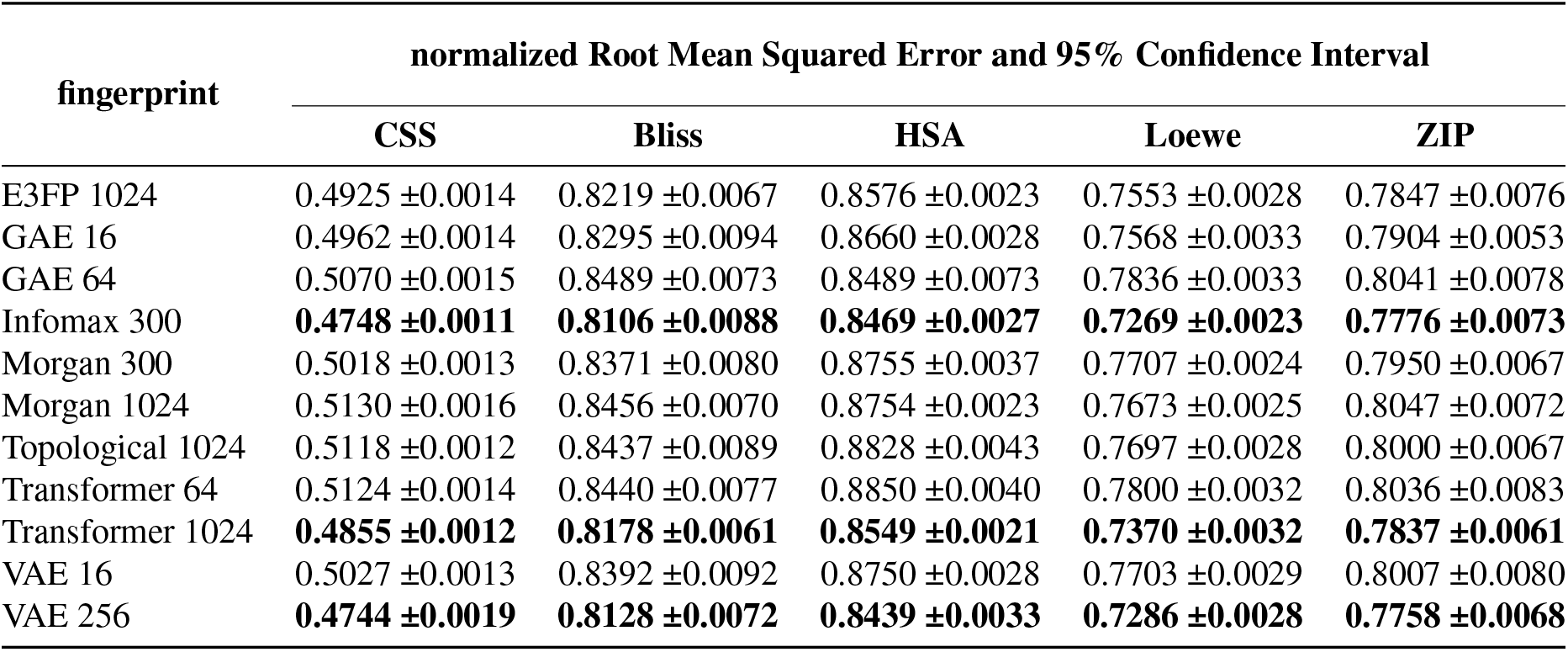
Drug combination synergy prediction on the CID-filtered dataset in 60 : 40 train:test split using RMSE, normalized by the target’s standard deviation. 95% confidence intervals are calculated via empirical bootstrap. Best models are highlighted with bold. VS I

| fingerprint | normalized Root Mean Squared Error and 95% Confidence Interval |  |  |  |  |
| --- | --- | --- | --- | --- | --- |
|  | CSS | Bliss | HSA | Loewe | ZIP |
| E3FP 1024 | 0.4925 ±0.0014 | 0.8219 ±0.0067 | 0.8576 ±0.0023 | 0.7553 ±0.0028 | 0.7847 ±0.0076 |
| GAE 16 | 0.4962 ±0.0014 | 0.8295 ±0.0094 | 0.8660 ±0.0028 | 0.7568 ±0.0033 | 0.7904 ±0.0053 |
| GAE 64 | 0.5070 ±0.0015 | 0.8489 ±0.0073 | 0.8489 ±0.0073 | 0.7836 ±0.0033 | 0.8041 ±0.0078 |
| Infomax 300 | **0.4748 ±0.0011** | **0.8106 ±0.0088** | **0.8469 ±0.0027** | **0.7269 ±0.0023** | **0.7776 ±0.0073** |
| Morgan 300 | 0.5018 ±0.0013 | 0.8371 ±0.0080 | 0.8755 ±0.0037 | 0.7707 ±0.0024 | 0.7950 ±0.0067 |
| Morgan 1024 | 0.5130 ±0.0016 | 0.8456 ±0.0070 | 0.8754 ±0.0023 | 0.7673 ±0.0025 | 0.8047 ±0.0072 |
| Topological 1024 | 0.5118 ±0.0012 | 0.8437 ±0.0089 | 0.8828 ±0.0043 | 0.7697 ±0.0028 | 0.8000 ±0.0067 |
| Transformer 64 | 0.5124 ±0.0014 | 0.8440 ±0.0077 | 0.8850 ±0.0040 | 0.7800 ±0.0032 | 0.8036 ±0.0083 |
| Transformer 1024 | **0.4855 ±0.0012** | **0.8178 ±0.0061** | **0.8549 ±0.0021** | **0.7370 ±0.0032** | **0.7837 ±0.0061** |
| VAE 16 | 0.5027 ±0.0013 | 0.8392 ±0.0092 | 0.8750 ±0.0028 | 0.7703 ±0.0029 | 0.8007 ±0.0080 |
| VAE 256 | **0.4744 ±0.0019** | **0.8128 ±0.0072** | **0.8439 ±0.0033** | **0.7286 ±0.0028** | **0.7758 ±0.0068** |

**Table S5:** Drug combination synergy prediction on the SMILES-filtered dataset in 90 : 10 train:test split using RMSE, normalized by the target’s standard deviation. 95% confidence intervals are calculated via empirical bootstrap. Best models are highlighted with bold. VS I

| fingerprint | normalized Root Mean Squared Error and 95% Confidence Interval |  |  |  |  |
| --- | --- | --- | --- | --- | --- |
|  | CSS | Bliss | HSA | Loewe | ZIP |
| E3FP 1024 | 0.4881 ±0.0033 | **0.7906 ±0.0148** | **0.8089 ±0.0131** | 0.7430 ±0.0080 | **0.7638 ±0.0162** |
| GAE 16 | 0.5041 ±0.0041 | 0.8203 ±0.0150 | 0.8351 ±0.0137 | 0.7788 ±0.0098 | 0.7845 ±0.0154 |
| GAE 64 | **0.4842 ±0.0037** | 0.7914 ±0.0182 | 0.8090 ±0.0136 | 0.7337 ±0.0084 | 0.7694 ±0.0193 |
| Infomax 300 | **0.4708 ±0.0038** | **0.7759 ±0.0155** | **0.7963 ±0.0133** | **0.7136 ±0.0092** | **0.7574 ±0.0182** |
| Morgan 300 | 0.4935 ±0.0033 | 0.8013 ±0.0153 | 0.8186 ±0.0132 | 0.7527 ±0.0083 | 0.7709 ±0.0148 |
| Morgan 1024 | 0.5045 ±0.0039 | 0.8077 ±0.0178 | 0.8242 ±0.0122 | 0.7541 ±0.0087 | 0.7793 ±0.0172 |
| Topological 1024 | 0.5276 ±0.0033 | 0.8122 ±0.0150 | 0.8294 ±0.0146 | 0.7530 ±0.0084 | 0.7889 ±0.0142 |
| Transformer 64 | 0.4972 ±0.0034 | 0.8082 ±0.0167 | 0.8222 ±0.0142 | 0.7545 ±0.0082 | 0.7776 ±0.0157 |
| Transformer 1024 | 0.4864 ±0.0033 | 0.7974 ±0.0157 | 0.8119 ±0.0149 | **0.7269 ±0.0084** | 0.7693 ±0.0150 |
| VAE 16 | 0.4919 ±0.0036 | 0.8034 ±0.0141 | 0.8174 ±0.0131 | 0.7509 ±0.0091 | 0.7737 ±0.0163 |
| VAE 256 | **0.4701 ±0.0037** | **0.7805 ±0.0162** | **0.7952 ±0.0146** | **0.7159 ±0.0080** | **0.7561 ±0.0159** |

**Table S6:**
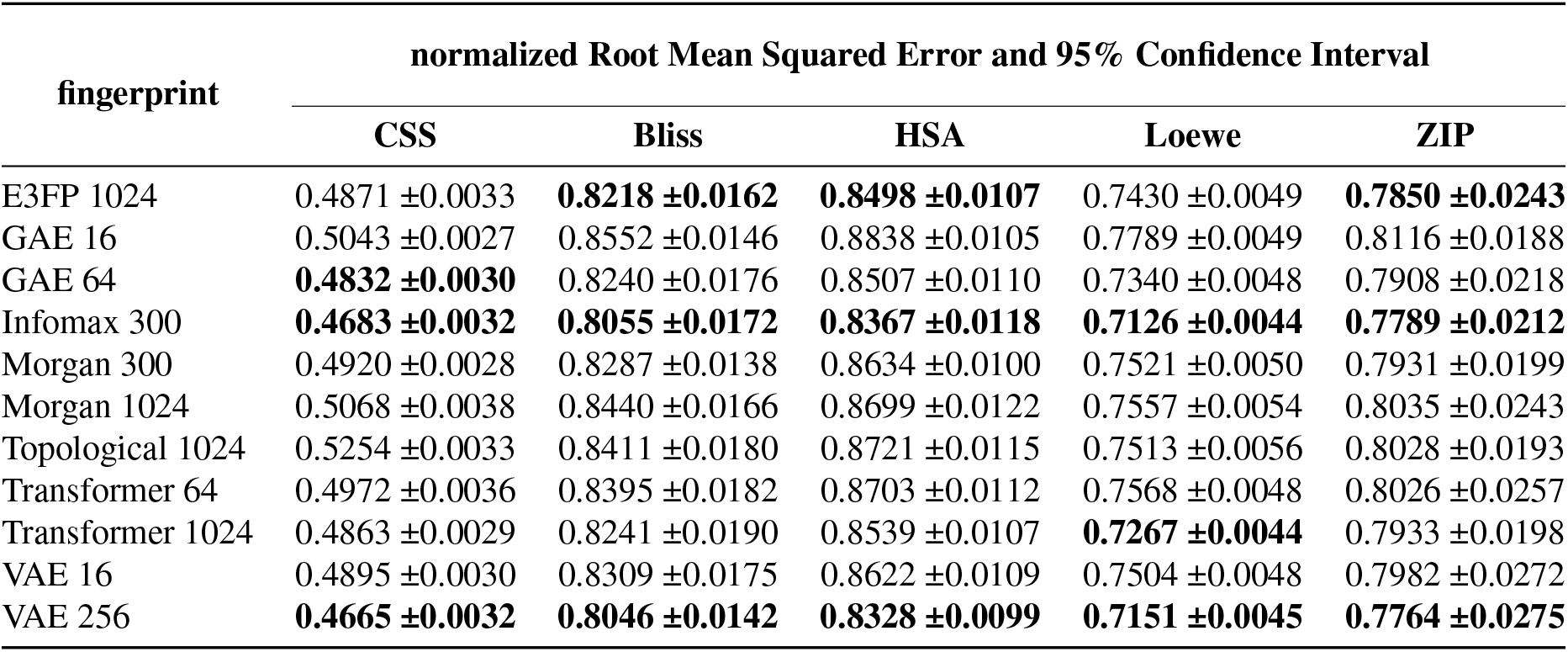
Drug combination synergy prediction on the CID-filtered dataset in 90 : 10 train:test split using RMSE, normalized by the target’s standard deviation. 95% confidence intervals are calculated via empirical bootstrap. Best models are highlighted with bold. VS I

## Notes

https://doi.org/10.5281/zenodo.4843919

### Competing Interest Statement

The authors have declared no competing interest.

### Summary of Updates

revision following reviewers' comments. Methods restructured for clarity; synergy models' description and prior work moved to Supplementary; fingerprint taxonomy table added. Style updates

